# BRIDGE: A Multi-organ Histo-ST Foundation Model Enables Virtual Spatial Transcriptomics for Enhanced Few-shot Cancer Diagnosis

**DOI:** 10.64898/2026.05.05.722971

**Authors:** Zhuo Liang, Weiqin Zhao, Fuying Wang, Guangyong Chen, Yuanhua Huang, Lequan Yu

## Abstract

Recent studies have explored generating virtual spatial transcriptomics (ST) profiles from histological images, offering a promising alternative to laboratory-measured molecular profiling. However, existing approaches predominantly rely on single-organ models and require substantial organ-specific training data, restricting their accuracy under challenging few-shot conditions in clincical practice, where less than 10 slides are available for specific organs or techniques. Here, we present BRIDGE, a multi-organ foundation model pre-trained on over 600,000 paired histology-ST profiles across 13 human organs and three sequencing techniques. By robustly aligning morphological features and genomic information within a shared multi-organ latent space, BRIDGE can leverage common biological knowledge across distinct tissues to enable accurate and generalizable pan-cancer molecular profiling. Without additional organ-specific fine-tuning, BRIDGE accurately predicts the spatial expression of 80 biomarker genes, achieving an average Pearson correlation coefficient (PCC) of 0.474—a 30% improvement over existing state-of-the-art models under three clinically challenging few-shot scenarios. With generated virtual ST, BRIDGE outperforms current state-of-the-art pathology foundation models in predicting cancer survival, achieving an average concordance index (C-index) of 0.724 across six TCGA cohorts. Notably, BRIDGE maintains exceptional performance even in zero-shot scenarios involving three cancer types not seen during its training, achieving an average C-index of 0.717, thereby demonstrating its strong generalization capability that transcends organ- and subtype-specific boundaries. Moreover, BRIDGE-generated virtual spatial transcriptomes match the prognostic accuracy of bulk RNA-seq, highlighting their potential as a spatially informative alternative to laboratory sequencing. In general, BRIDGE represents a data-efficient tool in virtual ST that facilitates biomedical discoveries in clinical few-shot contexts and advances diagnosis of understudied cancers without sufficient samples.

## INTRODUCTION

Histological imaging provides detailed morphological information crucial for clinical diagnostics, yet it lacks molecular resolution essential for understanding cancer biology and disease mechanisms. Spatial transcriptomics (ST) addresses this limitation by enabling gene expression analysis within the spatial context of pathological sections^1^. However, the high cost, technical complexity, and privacy concerns associated with generating ST data severely limit the availability of large-scale datasets. Thus, computationally integrating histological imaging with spatial transcriptomics to infer virtual profiling directly from routine histology slides presents a promising, scalable, and cost-effective alternative for cancer diagnosis (Fig. 1a).

**Figure 1.**
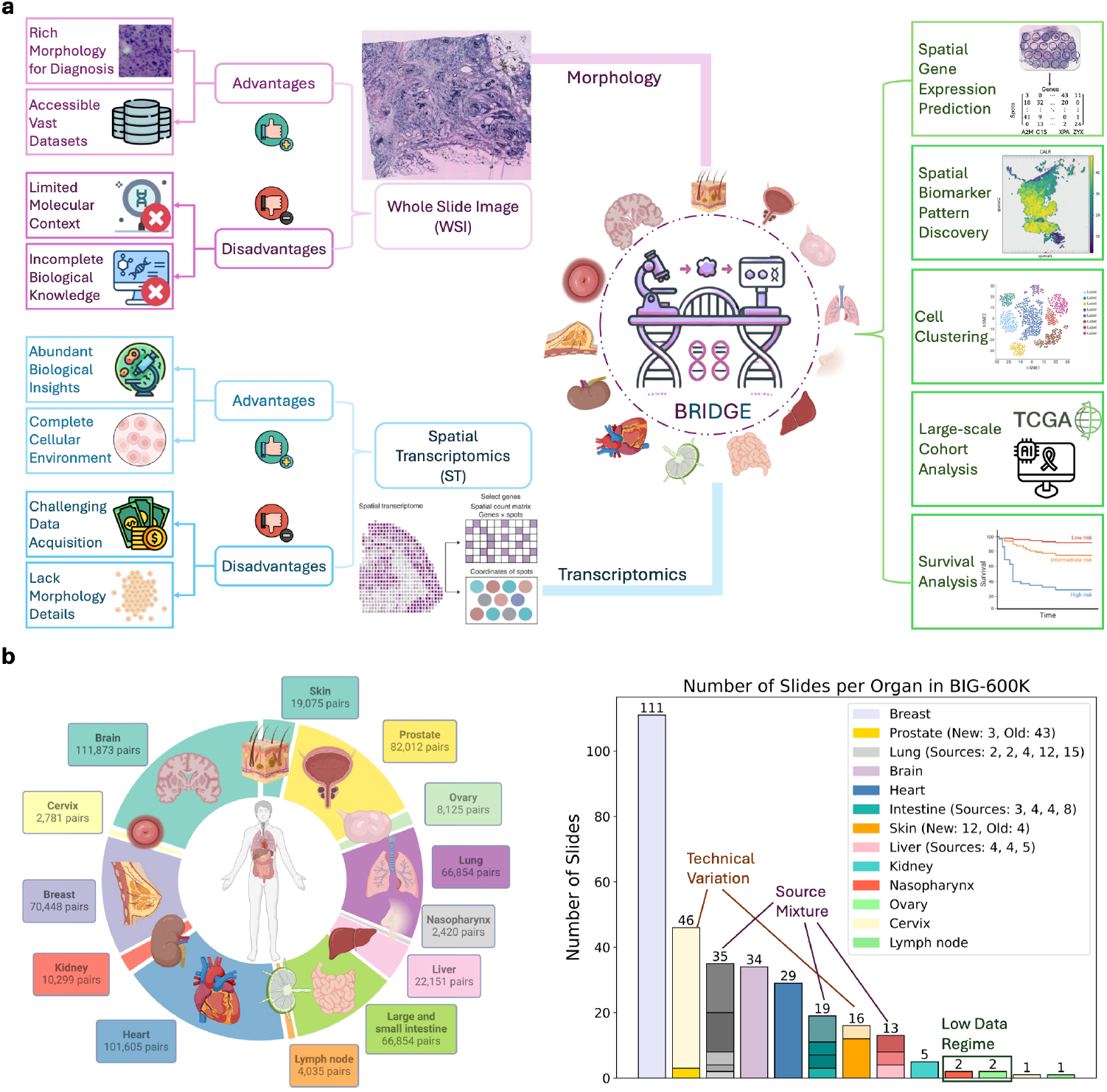
Rationale for integrating histological imaging with spatial transcriptomics (ST) and overview of the BIG-600K dataset. **a**, Histology images and spatial transcriptomics (ST) exhibit complementary strengths and limitations: histology provides detailed tissue morphology, whereas ST offers spatially resolved gene expression profiles. Integrating these modalities across multiple organs leverages shared biological knowledge, enabling more robust and comprehensive insights into tissue biology. **b**, Summary of the BIG-600K dataset, comprising over 600,000 paired histology-ST profiles across 13 human organs. The bar plot illustrates the distribution of slides per organ, highlighting dataset heterogeneity essential for robust multi-organ training of BRIDGE. We emphasize three challenging few-shot scenarios commonly encountered in ST data: (1) Low-Data Regime: Under-investigated organs (e.g., nasopharynx, ovary) with extremely limited available data (only two slides from two donors). (2) Technical Variation: Organs (e.g., prostate, skin) exhibiting significant imbalance between newer (e.g., 10X) and older techniques (e.g., ST), resulting in resolution differences of up to tenfold (approximately 3,000 vs. 300 spots per slide). (3) Source Mixture: Organs (e.g., lung, intestine, liver) integrating data from multiple laboratories, causing amplified batch effects due to varying instrument precision and protocols.

Recent deep learning approaches have attempted to bridge this modality gap between histology and ST. For instance, ST-Net^2^ predicted spatial expression of 250 target genes from histological images using a convolutional neural network pre-trained on natural images (e.g., ImageNet) combined with a fully connected layer as prediction head. Subsequent models^3–9^, including HE2RNA^3^, EMO^4^, HisToGene^5^, and DeepSpaCE^6^, extended this pipeline by investigating alternative image encoders and gene prediction heads. Graph-based methods like Hist2ST^10^, TransformerST^11^, TCGN^12^, and THItoGene^13^ further improved predictive accuracy by explicitly modeling spatial cell organization, while contrastive learning approaches such as BLEEP^14^ refined the integration of histological morphology and transcriptomic data. Despite these advances, nearly all existing methods rely on single-organ training strategies, which require substantial organ-specific data to avoid overfitting (Supplementary Table S1). Consequently, these organ-specific models exhibit limited generalizability, particularly under clinically realistic few-shot scenarios—characterized by minimal data availability, severe technical variability, and significant batch effects arising from multiple laboratory sources (Fig. 1b). Without adequate and diverse training data, existing methods frequently yield negative Pearson correlation coefficients (PCC) and fail to capture biologically meaningful spatial gene-expression patterns relevant to clinical diagnostics.

To systematically overcome these limitations, we constructed **BIG-600K** (**B**i-modal Dataset for Histology **I**maging and **G**ene Expression), comprising over 600,000 rigorously curated histology-ST pairs across 13 diverse human organs (Fig. 1b; Supplementary Table S2). BRIDGE integrates generative and contrastive learning strategies to robustly align morphological features from histological images with corresponding genomic information from spatial transcriptomics within a shared multi-organ latent space, thus enabling accurate and generalizable pan-cancer molecular profiling by effectively leveraging common biological knowledge across distinct tissues.

Without requiring organ-specific fine-tuning, BRIDGE accurately predicts spatial expression for 80 biomarker genes critical for cancer diagnostics and hereditary-disease identification. Under clinically realistic few-shot conditions, BRIDGE achieves an average PCC of 0.474—representing a 30% improvement over state-of-the-art single-organ methods. Beyond direct predictive inference, BRIDGE uniquely enables retrieval-based inference, matching histological images to laboratory-measured genomic profiles in large-scale sequencing reference databases. BRIDGE consistently outperforms existing retrieval methods by at least 60% in PCC across various retrieval scenarios and enables the generation of high-resolution spatially resolved virtual profiles derived from scRNA-seq data. By explicitly integrating genomic knowledge, BRIDGE significantly outperforms current state-of-the-art pathology foundation models (e.g., UNI) in cancer survival prediction across six TCGA cohorts. Remarkably, BRIDGE further demonstrates exceptional zero-shot generalization to three cancer types (bladder, gastric, esophageal) unseen during training, achieving an average C-index of 0.717. BRIDGE-generated virtual spatial transcriptomes match or surpass the prognostic performance of experimentally measured bulk RNA sequencing across multiple cancer types, including challenging zero-shot scenarios. Unlike bulk RNA-seq—which is costly and inherently lacks spatial context—BRIDGE provides biologically rich, spatially resolved molecular information directly from readily available clinical histology slides, thereby substantially reducing barriers to clinical adoption and facilitating deeper insights into tumor biology. In summary, BRIDGE represents a data-efficient computational framework that generates virtual spatial transcriptomics given routine histology slides from various organs and sequencing platforms, advancing diagnostic accuracy—particularly for understudied cancers lacking sufficient data.

## RESULTS

We provide a schematic overview of the BRIDGE framework in Fig. 2. BRIDGE integrates generative and contrastive learning strategies to support two complementary spatial transcriptomics inference tasks: direct prediction and retrieval-based inference, each offering distinct merits. Specifically, BRIDGE employs three interconnected learning objectives during training: (1) a contrastive loss, aligning visual and genomic modalities into a unified latent representation; (2) a generative loss, directly enhancing gene expression prediction accuracy from histological morphology; and (3) a reconstruction loss, preserving genomic information fidelity within the shared latent representation. For prediction tasks, BRIDGE leverages its pretrained histology image encoder and dedicated image-to-gene prediction module to infer spatially resolved gene expression profiles directly from previously unseen histological images. For retrieval tasks, BRIDGE utilizes reference gene expression datasets—multi-organ ST, single-organ ST, or single-cell RNA sequencing (scRNA-seq)—to identify and retrieve the most similar genomic profiles based on morphological features extracted from query histology images. To further demonstrate BRIDGE’s biological and clinical utility, we apply these inferred spatial gene expression profiles to cancer survival analyses using large-scale histological images from The Cancer Genome Atlas (TCGA), illustrating BRIDGE’s potential for precision cancer prognosis in clinical settings.

**Figure 2.**
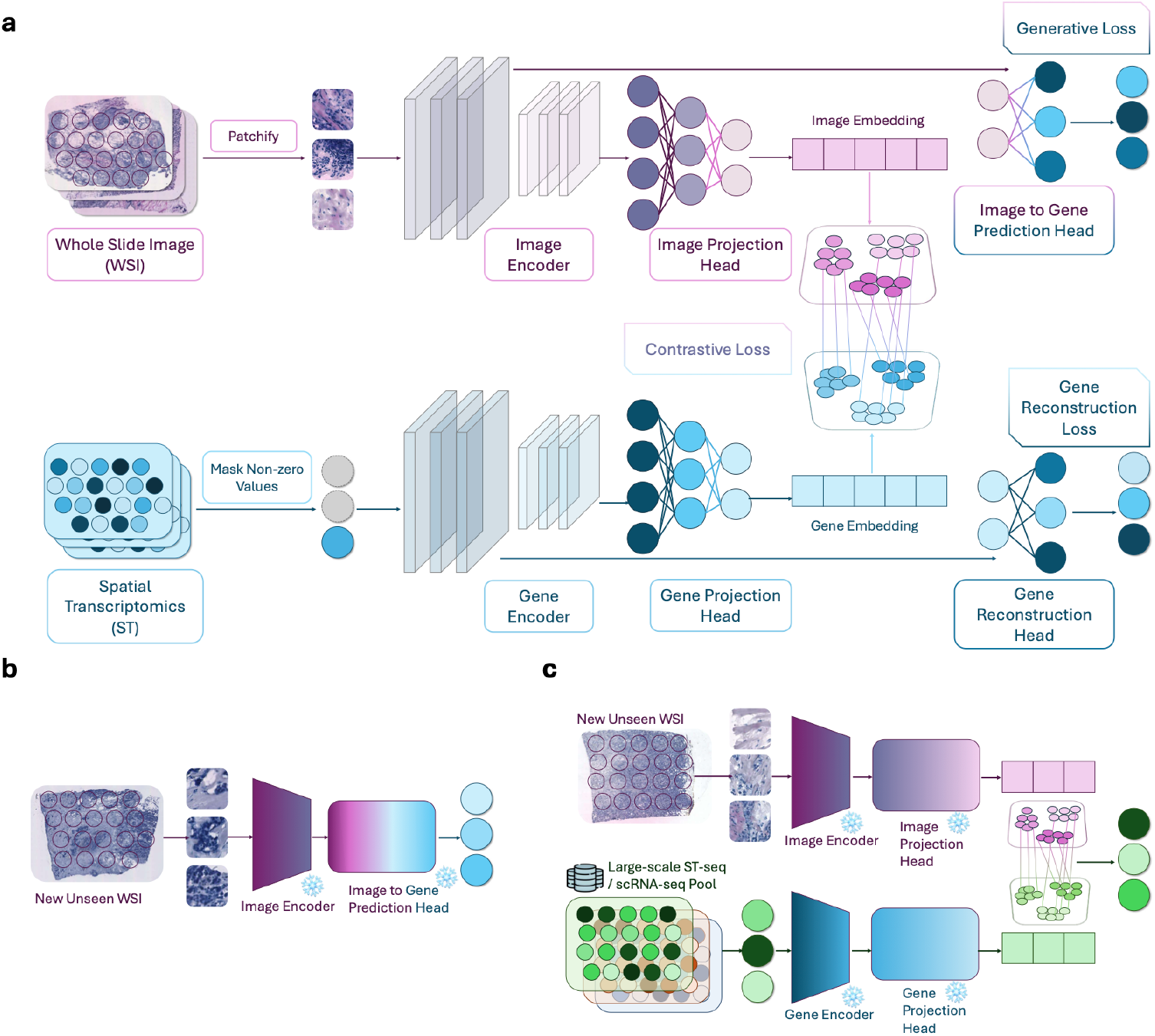
Overview of the BRIDGE framework. **a**, Schematic illustration of BRIDGE’s bi-modal architecture. BRIDGE simultaneously optimizes three complementary objectives—contrastive, generative, and reconstruction losses—to effectively align morphological (image-based) and genomic (gene-based) modalities into a shared latent representation space. **b**, Illustration of BRIDGE’s zero-shot spatial gene expression prediction task, where BRIDGE directly infers gene expression profiles from previously unseen histological images using its pretrained histology image encoder and dedicated image-to-gene prediction module. **c**, Illustration of BRIDGE’s retrieval-based inference task, applicable when suitable reference gene expression data (e.g., multi-organ ST, single-organ ST, or scRNA-seq) are available. BRIDGE retrieves the most similar gene expression profiles from the reference pool by matching histological image features within the shared latent representation space.

### BRIDGE Accurately Predicts Spatial Expression of Biomarker Genes Under Clinically Relevant Few-shot Conditions

We first evaluated BRIDGE’s predictive performance in inferring biomarker gene expression across seven diverse human organs under three clinically relevant few-shot scenarios frequently encountered in biomedical research and clinical practice: (1) low-data availability (understudied organs), (2) technical variation (imbalanced sequencing platforms), and (3) source mixture (multiple laboratories) (Fig. 1b, Fig. 3). The selected organs—ovary, nasopharynx, skin, prostate, lung, intestine, and liver—represent diverse biological contexts and clinical relevance. Our evaluation specifically focused on 80 biomarker genes critical for cancer diagnostics and hereditary-disease identification (Supplementary Table S3). We compared BRIDGE against several state-of-the-art (SOTA) methods, including ST-Net [2], DeepSpaCE [6], and graph-based models such as Hist2ST [10] and THItoGene [13].

**Figure 3.**
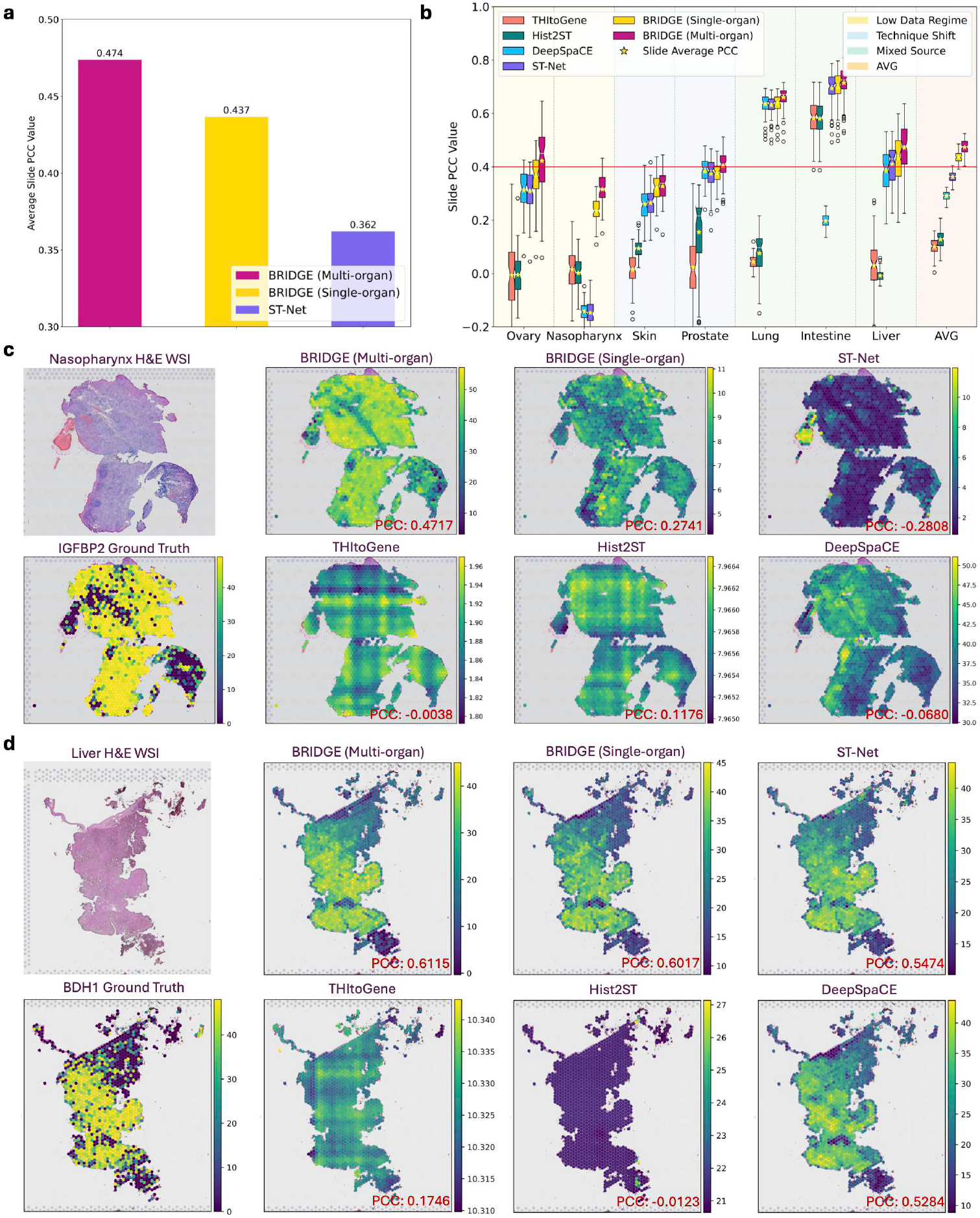
BRIDGE significantly improves spatial gene expression prediction across clinically challenging few-shot scenarios. **a**, Performance comparison between BRIDGE and previous SOTA baseline ST-Net under single-organ and multi-organ training settings. BRIDGE surpasses ST-Net in both scenarios, achieving average slide-level PCC of 0.437 (single-organ) and 0.474 (multi-organ), compared to ST-Net’s 0.362. **b**, Detailed performance evaluation across multiple baseline methods and BRIDGE under three clinically relevant few-shot scenarios: low-data availability (ovary, nasopharynx), technical variation (skin, prostate), and source mixture (lung, intestine, liver). AVG denotes average PCC across seven evaluated organs. **c**, Spatial prediction visualization for nasopharynx biomarker IGFBP2 under extreme low-data conditions (one training slide only). BRIDGE accurately reconstructs true expression patterns, outperforming baselines. **d**, Spatial prediction visualization for liver biomarker BDH1. BRIDGE consistently achieves highest PCC, clearly outperforming competing methods and accurately reconstructing true spatial gene expression patterns.

#### Multi-organ BRIDGE Outperforms Existing Single-organ Spatial Transcriptomics Methods

We initially compared our single-organ BRIDGE model (trained separately on each organ’s subset of BIG-600K) against ST-Net, previously the best-performing baseline method. As illustrated in Fig. 3a, single-organ BRIDGE significantly outperformed ST-Net, achieving an average Pearson correlation coefficient (PCC) of 0.437 compared to ST-Net’s 0.362. This improvement can be attributed primarily to BRIDGE’s explicit integration of contrastive and generative learning objectives, enabling more effective capture of cross-modality relationships between histological morphology and genomic features. Importantly, the multi-organ BRIDGE model—trained jointly across all 13 organs in BIG-600K—further improved the prediction accuracy to an average slide-level PCC of 0.474. This performance gain highlights the significance of heterogeneous multi-organ pretraining, allowing BRIDGE to learn generalized biological features robustly transferable across diverse organs and sequencing platforms. Individual slide-level analyses (Fig. 3b) and comprehensive evaluations across ten organs (including three non-few-shot organs: brain, breast, heart; Fig. S1)) consistently demonstrated BRIDGE’s superior performance, emphasizing the clinical value of multi-organ joint learning.

#### BRIDGE Demonstrates Robustness Under Extreme Low-data Conditions

Data scarcity poses a severe challenge for spatial transcriptomics prediction, particularly evident in understudied cancers. For example, BIG-600K includes only two slides each for ovary and nasopharynx (from different donors). With one slide used for training and the other for testing, substantial donor-specific variability introduced significant batch effects, negatively impacting baseline methods: graph-based models yielded negative PCC for ovary, while DeepSpaCE and ST-Net produced negative PCC for nasopharynx (Fig. 3b). Despite these difficulties, multi-organ BRIDGE consistently delivered robust predictions (PCC > 0.30) even under these stringent low-data conditions. Such robustness arises from BRIDGE’s effective leveraging of shared biological knowledge across multiple tissues learned during heterogeneous multi-organ training.

#### BRIDGE Effectively Mitigates Batch Effects from Technical Variation and Multiple Laboratories

Another critical challenge arises from batch effects due to differences in sequencing technologies and laboratory conditions. For instance, the skin dataset was imbalanced, comprising 25% older Spatial Transcriptomics (ST) slides (low sampling resolution) and 75% newer 10X Visium slides (higher sampling resolution). Prostate exhibited the opposite imbalance (93.5% older ST vs. 6.5% newer 10X). Consequently, baseline methods struggled to accurately predict gene expression on minority-platform slides (Fig. 3b). Similarly, lung, intestine, and liver datasets combined multiple laboratory sources (five, four, and three sources, respectively), amplifying batch effects from varying donor populations, instruments, and protocols. Remarkably, multi-organ BRIDGE effectively addressed these batch effects by leveraging mutual biological and technical insights from other organs sharing similar characteristics, consistently outperforming baseline methods and demonstrating strong generalizability across realistic clinical scenarios.

#### BRIDGE Faithfully Captures Spatial Patterns of Clinically Relevant Biomarker Genes

We further assessed BRIDGE’s capability to reconstruct spatially resolved gene expression patterns by visualizing predictions for two exemplary biomarker genes. In the challenging low-data scenario (nasopharynx biomarker IGFBP2 [15]; Fig. 3c), baseline methods either produced divergent spatial distributions with negative PCC (ST-Net, THItoGene, DeepSpaCE) or failed to accurately capture expression magnitude (Hist2ST). In contrast, BRIDGE achieved a gene-wise PCC of 0.472, closely matching ground-truth expression magnitude and spatial localization. Likewise, in a scenario with more available data (liver biomarker BDH1 [16, 17]; Fig. 3d), BRIDGE maintained clear superiority (PCC: 0.612), precisely reconstructing the true spatial gene expression pattern. These analyses demonstrate BRIDGE’s strength in capturing biologically relevant spatial transcriptomic structures accurately.

### BRIDGE Effectively Retrieves High-fidelity Spatially Resolved Gene Expression Profiles Across Diverse Retrieval Scenarios

In addition to direct prediction, BRIDGE supports retrieval-based inference of spatial gene expression by matching histological images to similar genomic profiles in its aligned latent space (Fig. 2c). Compared with direct prediction, retrieval-based inference can better preserve biologically meaningful gene-gene relationships by leveraging existing high-quality transcriptomic reference datasets. Furthermore, aggregating retrieved gene expression data from diverse sources can mitigate batch effects and discrepancies introduced by variability in tissue origins, donor populations, and sequencing platforms (Fig. 4a). To comprehensively assess BRIDGE’s retrieval capabilities, we compared it with BLEEP [14], a state-of-the-art retrieval-based spatial transcriptomics imputation method. Unlike BLEEP—which relies on simplistic retrieval pools composed of consecutive tissue sections from a single donor—we systematically evaluated BRIDGE across biologically and clinically relevant retrieval configurations. Specifically, we defined retrieval pools based on three critical dimensions: (1) organ specificity (single-organ vs. multi-organ), (2) sequencing modality (ST-seq vs. scRNA-seq), and (3) patient heterogeneity (mixed-patient vs. donor-paired), resulting in four distinct configurations (Fig. 4a). Due to current data unavailability, the “Single-organ Paired-patient scRNA-seq Pool” requiring paired scRNA-seq and ST-seq data from identical donors) remains a promising future direction.

**Figure 4.**
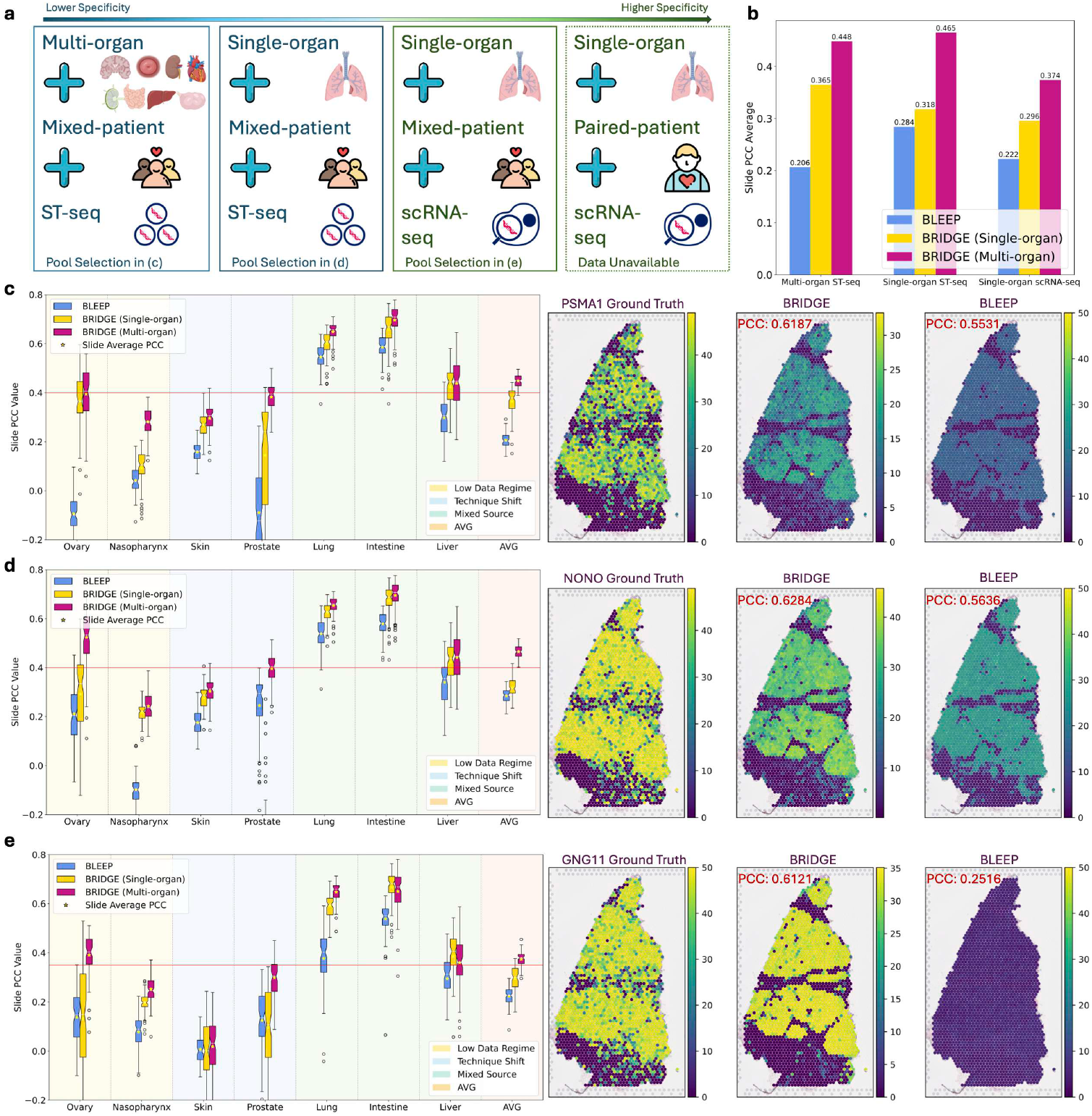
BRIDGE consistently retrieves high-fidelity spatial gene expression profiles across diverse retrieval pool configurations, substantially outperforming the state-of-the-art retrieval model BLEEP. **a**, Schematic overview of four biologically and clinically meaningful retrieval pool configurations, arranged from lower to higher specificity: Multi-organ Mixed-patient ST-seq Pool, Single-organ Mixed-patient ST-seq Pool, Single-organ Mixed-patient scRNA-seq Pool, and Single-organ Paired-patient scRNA-seq Pool (future direction due to current data limitations). These scenarios differ by organ specificity, sequencing modality, and donor heterogeneity. **b**, Average slide-level PCC comparison between BRIDGE and BLEEP across three evaluated retrieval pool configurations, demonstrating BRIDGE’s consistent superior retrieval performance (relative improvement ≥60%). **c**, Retrieval performance using Multi-organ Mixed-patient ST-seq pool (average slide PCC: BRIDGE 0.448 vs. BLEEP 0.206). Spatial visualization of lung cancer biomarker NONO highlights BRIDGE’s accurate spatial retrieval (gene-level PCC 0.619), significantly surpassing BLEEP. **d**, Retrieval results for Single-organ Mixed-patient ST-seq pool (average PCC: BRIDGE 0.465 vs. BLEEP 0.284). Visualization of biomarker PSMA1 confirms precise organ-specific retrieval by BRIDGE (gene-level PCC 0.628). **e**, Retrieval from Single-organ Mixed-patient scRNA-seq pool demonstrates BRIDGE’s robust cross-modality retrieval accuracy (average PCC: BRIDGE 0.374 vs. BLEEP 0.222), exemplified by accurate spatial prediction of biomarker gene GNG11 (gene-level PCC 0.612), while BLEEP fails to capture meaningful spatial patterns.

#### Organ-agnostic Retrieval Enables Robust Profiling Without Prior Information

A key clinical challenge is that the precise organ origin of tissue specimens may initially be unknown or ambiguous. Additionally, tissues across distinct organs often share similar cell types, biological functions, or disease-related gene expression signatures. Therefore, constructing a multi-organ retrieval pool is biologically meaningful, potentially enabling robust retrieval without requiring prior organ-specific knowledge. We first evaluated BRIDGE using the Multi-organ Mixed-patient ST-seq pool—representing the entire BIG-600K dataset encompassing diverse tissue origins and donor populations (Fig. 4b,c & Supplementary Fig. S4). BRIDGE consistently outperformed BLEEP, achieving an average slide-level PCC of 0.448 compared to 0.206 (BLEEP). Visualization of the biomarker gene NONO, associated with tumor progression and prognosis in lung cancer [18, 19], further highlighted BRIDGE’s superior spatial retrieval accuracy, achieving a gene-level PCC exceeding 0.61 and significantly surpassing BLEEP (Fig. 4c & Supplementary Fig. S6). These results underscore BRIDGE’s capacity to leverage crossorgan biological similarities, facilitating reliable, organ-agnostic spatial gene expression inference broadly applicable in clinical contexts.

#### Organ-agnostic Retrieval Enables Robust Profiling Without Prior Information

A key clinical challenge is that the precise organ origin of tissue specimens may initially be unknown or ambiguous. Additionally, tissues across distinct organs often share similar cell types, biological functions, or disease-related gene expression signatures. Therefore, constructing a multi-organ retrieval pool is biologically meaningful, potentially enabling robust retrieval without requiring prior organ-specific knowledge. We first evaluated BRIDGE using the Multi-organ Mixed-patient ST-seq pool—representing the entire BIG-600K dataset encompassing diverse tissue origins and donor populations (Fig. 4b,c & Supplementary Fig. S4). BRIDGE consistently outperformed BLEEP, achieving an average slide-level PCC of 0.448 compared to 0.206 (BLEEP). Visualization of the biomarker gene NONO, associated with tumor progression and prognosis in lung cancer [18, 19], further highlighted BRIDGE’s superior spatial retrieval accuracy, achieving a gene-level PCC exceeding 0.61 and significantly surpassing BLEEP (Fig. 4c & Supplementary Fig. S6). These results underscore BRIDGE’s capacity to leverage crossorgan biological similarities, facilitating reliable, organ-agnostic spatial gene expression inference broadly applicable in clinical contexts.

#### Organ-specific Retrieval Enables Precise Spatial Molecular Profiling

When organ-specific information is available, retrieval pools tailored to individual organs can deliver more concentrated biological and diagnostic molecular information. Unlike BLEEP—which requires transcriptome data from consecutive tissue sections of the same donor, severely restricting clinical applicability—BRIDGE imposes no such strict constraints. Specifically, BRIDGE leverages mixed-donor retrieval pools, significantly expanding practical utility, particularly in clinical scenarios where only a single slide per donor is typically available. We evaluated BRIDGE’s retrieval performance using Single-organ Mixed-patient ST-seq pools constructed from corresponding organ subsets within BIG-600K, focusing retrieval on organ-specific genomic profiles (Fig. 4b,d & Supplementary Fig. S4). BRIDGE demonstrated superior retrieval accuracy, attaining an average slide-level PCC of 0.465, substantially outperforming BLEEP (0.284). Spatial visualization of the biomarker gene PSMA1—clinically relevant across multiple cancers, including lung squamous cell carcinoma [20], hepatocellular carcinoma [21], and gastric cancer [22]—confirmed BRIDGE’s precise organ-specific molecular profiling capability (Fig. 4d & Supplementary Fig. S5). These results clearly highlight BRIDGE’s clinical utility to generate high-fidelity, organ-specific genomic profiles essential for targeted diagnostics and personalized therapeutic strategies.

#### BRIDGE Unlocks High-resolution Spatial Genomic Profiles by Bridging ST-seq and scRNA-seq Modalities

Single-cell RNA sequencing (scRNA-seq) typically provides genomic resolution at the cellular level superior to ST-seq, offering detailed biological insights at lower cost and greater accessibility given extensive public scRNA-seq datasets. Leveraging scRNA-seq data as a retrieval pool thus presents an exciting opportunity to generate high-resolution spatial genomic profiles. However, substantial technical and biological disparities between ST-seq and scRNA-seq traditionally limit effective cross-modality retrieval. Remarkably, BRIDGE successfully bridges these two modalities, effectively retrieving spatial gene expression profiles from Single-organ Mixed-patient scRNA-seq pools (Fig. 4b,e & Supplementary Fig. S4). Despite inherent differences between sequencing modalities, BRIDGE achieved robust retrieval performance, with an average slide-level PCC of 0.374, significantly outperforming BLEEP (0.222). Spatial visualization of the clinically significant biomarker gene GNG11 (lung adenocarcinoma [23], ovarian serous cystadenocarcinoma [24]) further illustrated BRIDGE’s impressive cross-modality retrieval accuracy (gene-level PCC: 0.612), whereas BLEEP failed to recapitulate meaningful spatial patterns (Fig. 4e). By effectively integrating transcriptomic information from both ST-seq and scRNA-seq modalities, BRIDGE unlocks novel clinical opportunities to generate cost-effective, high-resolution spatial transcriptomics.

### BRIDGE Delivers Accurate Cancer Prognosis and Robustly Generalizes in Zeroshot Scenarios

Accurate survival analysis—predicting critical clinical outcomes such as patient mortality and disease progression—is essential for personalized treatment planning, clinical decision-making, and effective patient risk stratification. [25] Here, we systematically evaluated BRIDGE’s ability to infer patient survival outcomes from routine clinical histology slides, using large-scale cancer cohorts from The Cancer Genome Atlas (TCGA) (Supplementary Table S4). Specifically, we assessed BRIDGE’s performance across six diverse cancer cohorts, including three cancer types unseen during training: Urothelial Bladder Carcinoma (BLCA), Esophageal Carcinoma (ESCA), and Stomach Adenocarcinoma (STAD). Additionally, we assessed BRIDGE on Breast Invasive Carcinoma (BRCA-HER2+ subtype), Triple-Negative Breast Cancer (BRCA-TNBC subtype, unseen subtype during training), and Lung Adenocarcinoma (LUAD). Notably, although BRCA-HER2+ and LUAD were included in training, the amount of training data represented is less than one-tenth of the corresponding TCGA testing cohorts, approximating a few-shot evaluation scenario. This rigorous evaluation allowed us to comprehensively investigate BRIDGE’s generalization capabilities under both zero-shot (unseen cancer types) and fewshot (limited training data for known cancer types) clinical contexts. We further benchmarked BRIDGE against (i) state-of-the-art pathology-image foundation models and (ii) conventional bulk RNA sequencing—the clinical reference for cancer diagnostics as it captures established diagnostic and prognostic gene signatures [26–32].

#### BRIDGE Achieves Robust Zero-shot Transfer To Unseen Cancer Types

A key challenge in clinical prognosis is accurately predicting outcomes for cancers absent from training datasets. Although BRIDGE was not trained on bladder, oesophageal, or gastric tumours, it nevertheless achieved competitive accuracy on these cancers. A plausible explanation is that the model transfers cross-organ transcriptional patterns that are partly conserved among anatomically or functionally related tissues—for example, kidney and bladder(anatomical and functional connections), intestine and oesophagus (joint digestive function), and liver, intestine and stomach (closely integrated roles in digestion and nutrient metabolism)—rather than relying solely on cancer-specific signals. BRIDGE achieved an impressive average concordance index (C-index) of 0.717 across these three unseen cancer cohorts (Table 1, Fig. 5b&c), clearly underscoring its robust zero-shot generalization capability. This finding significantly expands BRIDGE’s clinical utility, particularly for under-investigated cancers with limited available data.

**Table 1.** Comparison of BRIDGE with state-of-the-art pathology and visual foundation models and bulk RNA-seq for cancer survival prediction across six TCGA cancer cohorts. The six cohorts represent three clinically relevant and challenging evaluation scenarios: (1) Organ Zero-shot cohorts (STAD, ESCA, BLCA), where BRIDGE received no organ-specific training data; (2) Subtype Zero-shot cohort (BRCA-TNBC), where BRIDGE was trained only on other breast cancer subtypes but not TNBC; and (3) Slide Few-shot cohorts (BRCA-HER2+, LUAD), where BRIDGE’s training dataset size was significantly smaller (less than 10%) compared to corresponding test cohort sizes. BRIDGE achieved the highest concordance index (C-index; highlighted in **red**) in three cancer cohorts (STAD, BRCA-HER2+, LUAD) and second-best performance (highlighted in **blue**) in the remaining three cohorts, attaining the best overall average rank (1.5) across all models and scenarios. Importantly, BRIDGE maintained superior performance even under challenging zero-shot conditions (average rank 1.67), surpassing large-scale pathology foundation models explicitly trained on corresponding organs (e.g., UNI pretrained on extensive esophagogastric and bladder data). Experimentally measured bulk RNA-seq results (shown in **gray**) provided as a practical upper-bound reference. Models are ordered by their overall average rank across all cohorts.

| Dataset | BRIDGE | CTransPath | UNI | CONCH | QuiltNet | PLIP | DINOv2 | HIPT | Bulk-RNA |
| --- | --- | --- | --- | --- | --- | --- | --- | --- | --- |
| **Organ Zero-shot Cohorts** |  |  |  |  |  |  |  |  |  |
| STAD | **67.13** | 64.71 | 65.22 | 64.61 | **66.01** | 64.13 | 64.55 | 60.35 | 65.13 |
| ESCA | **71.44** | 70.94 | **71.64** | 68.61 | 68.42 | 67.55 | 67.84 | 63.43 | 69.50 |
| BLCA | **76.40** | **76.70** | 75.73 | 75.94 | 74.92 | 76.24 | 73.94 | 73.30 | 79.24 |
| **Subtype Zero-shot Cohort** |  |  |  |  |  |  |  |  |  |
| BRCA-TNBC | **75.13** | 74.27 | 74.35 | **78.27** | 74.57 | 69.68 | 72.74 | 71.57 | 70.60 |
| **Slide Few-shot Cohorts** |  |  |  |  |  |  |  |  |  |
| BRCA-HER2+ | **70.44** | **67.78** | 64.05 | 66.14 | 64.85 | 66.57 | 65.82 | 57.82 | 70.75 |
| LUAD | **73.73** | 70.90 | **71.57** | 69.63 | 70.40 | 70.55 | 70.78 | 71.02 | 75.44 |
| **Averages by Setting** |  |  |  |  |  |  |  |  |  |
| Organ Zero-shot (Avg. C-idx) | **71.66** | **70.78** | 70.86 | 69.72 | 69.78 | 69.31 | 68.78 | 65.69 | 71.29 |
| Organ Zero-shot (Avg. Rank) | **1.67** | **2.67** | 3.00 | 4.33 | 4.33 | 5.67 | 6.33 | 8 | N/A |
| All Cohorts (Avg. C-idx) | **72.38** | **70.88** | 70.43 | 70.53 | 69.86 | 69.12 | 69.28 | 66.25 | 71.78 |
| All Cohorts (Avg. Rank) | **1.5** | **3.17** | 3.67 | 4.33 | 4.83 | 5.67 | 5.83 | 7 | N/A |

**Figure 5.**
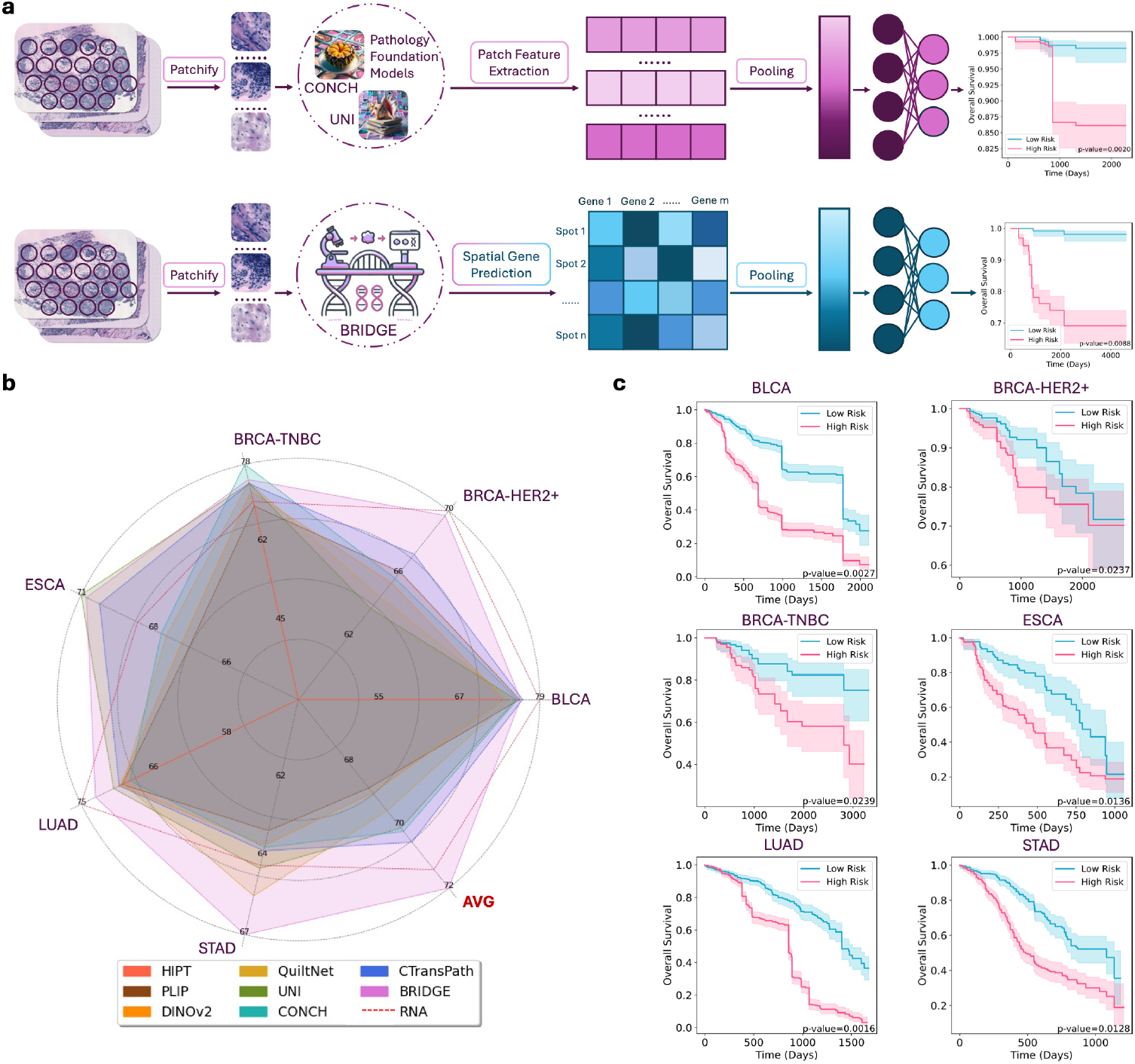
BRIDGE accurately predicts patient survival outcomes across multiple TCGA cancer cohorts, robustly generalizing to unseen cancer types (zero-shot), surpassing state-of-the-art pathology foundation models, and matching or exceeding bulk RNA-seq performance. **a**, Methodological comparison between BRIDGE and conventional pathology foundation models. Pathology models extract visual features from patches within whole-slide images (WSIs), aggregating patch-level visual embeddings into slide-level representations via averaging or attention pooling. In contrast, BRIDGE infers spatial gene-expression profiles at the spot-level from histological patches, aggregating these spot-level genomic predictions into bulk RNA-seq-like slide-level representations. Both approaches employ a multi-layer perceptron (MLP) with Cox regression to predict patient survival risk, optimizing MLP parameters during training. **b**, Radar chart comparing BRIDGE’s concordance index (C-index) with seven leading pathology foundation models across six cancer cohorts, including three completely unseen cancers (BLCA, ESCA, STAD). The red dotted line indicates bulk RNA-seq performance, typically considered the upper bound for genomic-based survival prediction. BRIDGE consistently demonstrates superior or comparable performance across all cohorts, clearly surpassing pathology models despite significantly fewer training samples and model parameters. (C-index scaled by 100 for visualization clarity.) **c**, Kaplan–Meier survival curves illustrating BRIDGE’s effective stratification of high- and low-risk patient groups across six cancers. Clear and statistically significant separation between risk groups highlights BRIDGE’s robust prognostic accuracy, even in challenging zero-shot scenarios.

#### BRIDGE Surpasses State-of-the-art Pathology Foundation Models Despite Smaller Pretraining Scale

We benchmarked BRIDGE against several leading pathology and visual foundation models, including CTransPath [33], UNI [34], CONCH [35], QuiltNet [36], PLIP [37], DINOv2 [38], and HIPT [39]. Notably, UNI and CONCH recently demonstrated state-of-the-art performance in clinical prognosis, benefiting from extensive pre-training on massive datasets. For instance, UNI was pretrained on over 100 million histology image patches from 20 organs, employing approximately 303 million parameters—vastly exceeding BRIDGE’s relatively modest ∼500K paired training samples and 19.6 million parameters. Despite this substantial difference in data scale and model complexity, BRIDGE consistently outperformed competing pathology foundation models, achieving the highest C-index in three cancer cohorts (STAD, BRCA-HER2+, LUAD) and second-best performance in the remaining three (ESCA, BLCA, BRCA-TNBC) (Fig. 5b, Table 1). Across all six cohorts, BRIDGE achieved an average rank of 1.5, clearly surpassing the next-best baseline (average rank of 3.17). Specifically, BRIDGE outperformed UNI by an average relative C-index improvement of 2.77% across all cohorts. Even in challenging zero-shot scenarios (STAD, ESCA, BLCA), BRIDGE outperformed UNI by a relative improvement of 1.13%, despite UNI’s explicit pretraining on 6,705 esophagogastric and 1,059 bladder cancer slides. These findings demonstrate BRIDGE’s superior clinical accuracy, computational efficiency, and accessibility compared with existing large-scale visual foundation models, significantly reducing barriers to clinical adoption.

#### BRIDGE Matches Bulk RNA-seq Performance, Offering a Cost-effective and Spatially Informative Alternative

Bulk RNA sequencing is widely regarded as the gold-standard method for transcriptomic-based clinical prognosis, directly quantifying gene expression averaged across cell populations. However, bulk RNA-seq is costly, frequently unavailable in clinical practice, and inherently lacks critical spatial context needed to understand tumor heterogeneity and microenvironment complexity. By accurately predicting spatially resolved gene-expression profiles from routine histology slides, BRIDGE captures molecular and spatial tumor characteristics simultaneously. Remarkably, BRIDGE matched or even surpassed bulk RNA-seq across six TCGA cohorts, achieving an average C-index of 0.724 compared with bulk RNA-seq’s 0.718 (Fig. 5b, Table 1). Notably, BRIDGE significantly outperformed bulk RNA-seq in challenging zero-shot settings for gastric (STAD; 0.671 vs. 0.651) and esophageal cancers (ESCA; 0.714 vs. 0.695). Moreover, BRIDGE demonstrated superior prognostic accuracy for rare but aggressive Triple-Negative Breast Cancer subtype (BRCA-TNBC), surpassing bulk RNA-seq by a relative margin of 6.42%. These results clearly highlight BRIDGE’s potential as a clinically accessible, cost-efficient, and spatially informative alternative to bulk RNA sequencing, significantly advancing precision diagnostics and personalized treatment planning, especially for understudied cancers.

## DISCUSSION

In this study, we introduce BRIDGE, a bi-modal multi-organ foundation model specifically designed for computational inference of spatially resolved gene expression directly from routine histological images. Leveraging our rigorously curated multi-organ dataset BIG-600K, BRIDGE robustly aligns morphological features with corresponding genomic information within a shared multi-organ latent representation. Unlike existing single-organ models—which require organ-specific training data of at least 10 routine slides—BRIDGE effectively leverages shared biological knowledge across diverse human tissues, enabling accurate inference even under few-shot and zero-shot conditions. By generating virtual spatial transcriptomics profiles directly from standard histology slides, BRIDGE provides a scalable, computationally efficient, and clinically accessible tool for cancer research.

Several recent multi-organ histology–ST datasets, such as HEST-1K [40] and STimage-1K4M [41], have provided valuable resources for biomedical research. However, the absence of foundational computational models accompanying these datasets has limited their practical clinical impact. In contrast, BRIDGE explicitly capitalizes on the substantial biological and technical heterogeneity captured within BIG-600K, directly translating these complexities into significantly improved generalization and robustness. Our systematic evaluation across multiple challenging few-shot scenarios—including low-data availability, pronounced technical variation in sequencing platforms, and significant batch effects arising from multi-laboratory datasets—consistently demonstrated BRIDGE’s marked superiority over existing single-organ approaches. These results underscore the critical importance of multi-organ pretraining for achieving robust generalization in virtual genomic profiling, highlighting BRIDGE’s potential as a scalable and reliable solution for future biomedical research and clinical diagnostics. A promising future direction involves fine-tuning BRIDGE’s prediction module with additional single-organ or cancer-specific datasets. Such targeted fine-tuning could further enhance predictive accuracy without sacrificing generalizability, potentially enabling even more precise diagnostic performance in clinical settings.

Beyond direct gene expression prediction, BRIDGE uniquely supports retrieval-based inference, inherently preserving biologically meaningful gene-gene relationships and mitigating batch effects and technical discrepancies. Our comprehensive retrieval experiments revealed that BRIDGE consistently outperformed existing state-of-the-art retrieval methods across diverse biologically relevant retrieval scenarios. Notably, BRIDGE effectively bridges spatial transcriptomics (ST-seq) and single-cell RNA sequencing (scRNA-seq) modalities, successfully retrieving high-resolution genomic profiles from large-scale scRNA-seq reference databases. Although current data limitations preclude evaluating donor-matched ST–scRNA-seq retrieval pools, future efforts to generate paired ST–scRNA-seq datasets could further enable precise genomic profiling for personalized clinical decision-making.

A particularly impactful finding is BRIDGE’s remarkable zero-shot generalization capability. BRIDGE successfully transferred genomic knowledge across anatomically and functionally related organs, accurately predicting survival outcomes for three cancer types unseen during training (bladder cancer [BLCA], esophageal carcinoma [ESCA], and stomach adenocarcinoma [STAD]). Furthermore, BRIDGE demonstrated robust subtype-level generalization, accurately predicting survival outcomes for the rare and aggressive triple-negative breast cancer subtype (BRCA-TNBC), despite never encountering this specific subtype during training. These findings strongly indicate BRIDGE’s powerful ability to learn generalizable molecular patterns that transcend organ- and subtype-specific boundaries, significantly expanding its clinical utility, particularly for understudied cancers lacking sufficient training data.

Despite its relatively modest model size (19.6 million parameters) and training dataset scale (∼500K paired samples), BRIDGE consistently outperformed leading pathology foundation models, including UNI and CONCH, which were pretrained on significantly larger datasets (>100 million histological patches and >300 million parameters for UNI). Importantly, even in zero-shot scenarios, BRIDGE outperformed UNI, despite UNI’s explicit pretraining on extensive esophagogastric and bladder cancer data. These compelling results highlight BRIDGE’s efficient bi-modal architecture, demonstrating that explicit genomic integration and multi-task training objectives substantially enhance predictive accuracy without requiring excessively largescale training data or computational resources. Finally, BRIDGE-generated virtual spatial transcriptomes matched or even exceeded the prognostic performance of bulk RNA sequencing— the widely considered gold standard for transcriptomic-based clinical prognosis—across multiple cancer cohorts. Unlike bulk RNA sequencing, which is costly, infrequently available in clinical practice, and inherently lacks spatial context, BRIDGE directly generates biologically rich, spatially informative molecular profiles from routine histology slides, providing deeper insights into tumor heterogeneity and microenvironmental complexity. BRIDGE thus represents a scalable, affordable, and broadly generalizable computational framework effectively bridging routine histopathology and precision genomics, significantly advancing diagnostic accuracy— particularly for clinically relevant yet underexplored cancers. With the continued growth of spatial transcriptomics technologies and datasets, we anticipate further improvements in BRIDGE’s performance and clinical utility, potentially reshaping cancer diagnostics toward affordable, spatially informed molecular profiling.

## METHODS

### Construction of the BIG-600K Dataset

In this section, we present detailed information on data sources, numerical compositions, and selection criteria employed during the construction of the BIG-600K dataset. Comprehensive statistics, including sampled organs, number of whole-slide images (WSIs), and spatial transcriptomic spots, are provided in Fig. 1b and Supplementary Table S2.

#### Data Sources

The BIG-600K dataset integrates spatial transcriptomics (ST) data from three primary sources: (1) publicly available experimental datasets gathered from peer-reviewed biomedical journals and preprint platforms such as bioRxiv (https://www.biorxiv.org/); (2) sequencing outputs from reputable biotechnology companies, notably 10x Genomics (https://www.10xgenomics.com/); and (3) open-access repositories including Dryad (https://datadryad.org/stash), Human Cell Atlas (https://www.humancellatlas.org/), Mendeley Data (https://data.mendeley.com/), and Zenodo (https://zenodo.org/). Relevant datasets were identified using keywords such as “spatial transcriptomics.” These sources were selected for their provision of high-quality, clinically relevant data suitable for robust evaluations and analyses.

#### Data Selection Criteria

We employed strict inclusion criteria to ensure data suitability: (1) samples derived exclusively from human tissues; (2) high-resolution histological images enabling precise delineation of cellular morphology; (3) comprehensive genomic coverage, typically involving thousands of genes measured per sample; and (4) precise lab-generated spatial coordinates, facilitating accurate alignment between histological features and spatial transcriptomic data with minimal manual adjustments.

### Data Preprocessing

In this section, we describe the complete preprocessing pipeline.

#### Histology Image Processing

To align histology images effectively with spatial transcriptomics data, we applied several preprocessing steps to whole-slide images (WSIs). Specifically, WSIs were cropped into patches sized 224 × 224 pixels, centered around lab-generated sequencing coordinates with a physical diameter of 150 *µ*m. Given the variability in spot sizes (55–100 *µ*m, sequencing method-dependent [42]), this cropping window ensured sufficient contextual information. Subsequently, image patches predominantly containing background areas were filtered out using instrument-provided tissue indicators and additional manual quality controls. To mitigate color variability resulting from batch effects, structure-preserving color normalization [43] was uniformly applied to all patches using a reference slide. Furthermore, various data augmentation techniques—including random cropping, flipping, rotation, and Gaussian blurring—were employed to enhance the robustness of the visual encoder within the bi-modal framework [44].

#### Spatial Transcriptome Data Processing

We standardized gene expression data across datasets by unifying gene identifiers. Given discrepancies in gene annotation formats (e.g., Human Ensembl gene IDs versus gene names), we constructed a comprehensive mapping dictionary using Visium-generated “features.tsv” files, supplemented by Ensembl’s BioMart^1^ when necessary. Ambiguous or deprecated gene IDs were excluded. Subsequently, we selected 7,730 genes consistently measured across multiple organs within BIG-600K, capturing essential molecular interactions while reducing dimensional complexity. To further reduce batch effects inherent in gene expression counts [45], we discretized raw counts into 50 bins following established preprocessing methodologies [46, 47]. Finally, we adopted a masked language modeling (MLM) approach [46, 47], randomly masking 25% of non-zero gene expression values to zero, enhancing the robustness of the genomic data encoder to sparse data.

#### Biomarker Gene Selection

From the pool of 7,730 cross-organ consistent genes, we further curated 80 biomarker genes with significant diagnostic implications for cancers and genetic disorders (Supplementary Table S3). The performance of BRIDGE and baseline methods on predicting these biomarkers is thoroughly evaluated in the Results section.

### Framework Architecture of BRIDGE

BRIDGE introduces a novel bi-modal architecture designed to overcome inherent limitations associated with conventional prediction-only or retrieval-only approaches for inferring spatial gene expression from histology images. Existing prediction-based methods often overlook complex molecular details extending beyond visual tissue features, resulting in predictions misaligned with biological measurements. Conversely, retrieval-based methods typically rely on rigid alignment between reference and query histology images, making them susceptible to domain shifts and technical variations. Moreover, previous retrieval approaches primarily use contrastive learning, potentially oversimplifying modality relationships by neglecting modality-specific information. BRIDGE addresses these limitations by integrating multiple learning objectives across distinct modules to effectively capture intra- and inter-modality relationships. Specifically, BRIDGE comprises three main components: Visual Encoder, Genome Encoder, and Gene Prediction and Reconstruction Decoder, each serving distinct roles in processing and aligning histological and genomic information.

#### Visual Encoder

We utilized a DenseNet-121 neural network [48], pre-trained on ImageNet [49], as the primary image encoder. Image patches I ∈ ℝ^B*×*3*×*224*×*224^ (with batch size B) were encoded into original and augmented feature vectors I_f_, 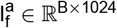. These vectors were projected into a shared 128-dimensional embedding space via a multilayer perceptron (MLP), generating image embeddings I_e_, 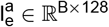.

#### Genome Encoder

For encoding gene expression data, we employed TabNet [50], a specialized architecture for tabular data. The input gene vector G ∈ ℝ^B*×*7730^ was masked randomly (25% of non-zero values set to zero), yielding G^m^ ∈ ℝ^B*×*7730^. TabNet encoded both original and masked gene vectors into genomic feature vectors G_f_, 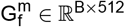, subsequently projected into the shared embedding space via another MLP, resulting in gene embeddings 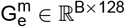.

#### Gene Prediction and Reconstruction Decoder

The gene prediction head employed fully connected layers to map visual features I_f_ directly to predicted gene expression profiles 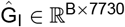. Concurrently, inspired by masked language modeling (MLM), the reconstruction head mapped masked genomic features 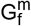 back to gene expression profiles 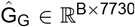, reinforcing the robustness of the genome encoder.

### Training Objectives of BRIDGE

BRIDGE integrates complementary learning objectives—including contrastive, generative, and reconstruction losses—to enhance overall modeling performance. These objectives enable effective modality alignment and robust feature encoding.

#### Contrastive Loss

To align visual and genomic embeddings, we applied contrastive learning via the InfoNCE loss [51] in two distinct alignment scenarios: (1) between original embeddings I_e_ and G_e_, and (2) between augmented image embeddings 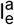 and masked gene embeddings 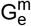. The two contrastive losses were computed as follows:

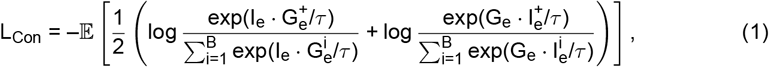

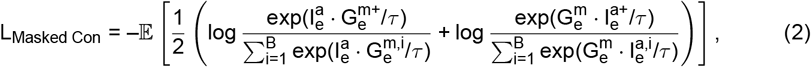

where the temperature parameter *τ* was set to 0.07, following prior literature [52].

#### Generative Loss

To enhance direct gene prediction from histology images and encourage the visual encoder to capture molecularly informative features, we introduced a generative loss based on Mean Squared Error (MSE):

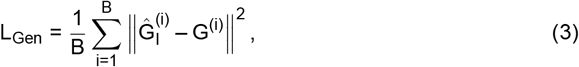

where 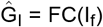 represents predictions from visual features.

#### Reconstruction Loss

We applied a reconstruction loss to ensure the genome encoder retained essential molecular information despite masking:

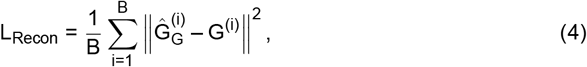

where 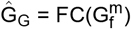 denotes reconstructed gene expressions from masked genomic features.

#### Modality-specific Supervision Loss

Additional modality-specific supervision losses were included to enhance intra-modality robustness [44]. For visual data, we employed SimCLR-based augmentation and contrastive learning [53]:

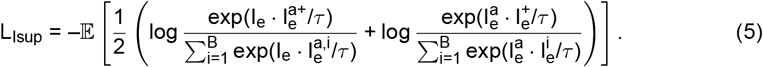

Similarly, for genomic data, we applied a contrastive loss between original and masked embeddings:

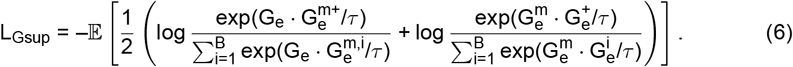

#### Overall Training Objective

The combined training objective integrated all loss functions with empirically determined weights:

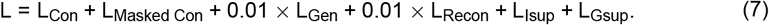

### Data Splits and Training Strategy

To comprehensively evaluate the performance of BRIDGE, we designed two distinct evaluation scenarios: (1) a challenging few-shot evaluation setting comprising seven WSIs from seven different organs, and (2) a more comprehensive multi-organ evaluation setting involving sixteen WSIs from ten human organs.

#### Few-shot Evaluation Setting

We reserved one WSI for testing from each of the seven organs characterized by challenging few-shot conditions: nasopharynx, ovary, skin, prostate, lung, intestine, and liver. This resulted in a total of 7 WSIs containing 22,526 image-gene pairs (spots).

#### Multi-organ Evaluation Setting

To further demonstrate BRIDGE’s generalizability across diverse organs and tissues, we also evaluated performance on a larger-scale multi-organ dataset encompassing ten human organs: brain, breast, heart, liver, lung, nasopharynx, ovary, prostate, skin, and intestine. For organs with more than 20 WSIs, two slides were reserved for testing; otherwise, one slide was reserved. This resulted in a comprehensive evaluation set consisting of 16 WSIs, comprising a total of 99,874 image-gene pairs (approximately 20% of the training dataset).

#### Single-Organ vs. Multi-Organ Training

We trained BRIDGE under two distinct configurations: single-organ training, where the model was trained individually for each organ, and multi-organ training, where training involved data from multiple organs simultaneously. Due to dataset size differences, the single-organ models converged after 20 epochs, whereas the multi-organ model (approximately 500K data pairs per epoch) converged after 50 epochs.

#### Technical Configurations

BRIDGE was trained using eight NVIDIA GeForce RTX 3090 GPUs with a total batch size of 384. We employed gradient accumulation (step size of 2), linear warmup, and cosine annealing learning rate schedules, with a peak learning rate of 1 × 10^−3^ for a maximum of 50 epochs.

### Application of BRIDGE on Downstream Tasks

In this section, we delineate how BRIDGE is applied to key downstream analyses, encompassing virtual gene prediction and retrieval as well as patient-level survival modelling.

#### Biomarker Gene Prediction and Retrieval

For the prediction task, pre-trained visual encoders and gene prediction modules directly inferred gene expressions from histology images. For retrieval, embeddings I_e_ and G_e_ were first generated from query histology images and reference genomic datasets. The Euclidean distance between query image embeddings I_e_ ∈ ℝ^B*×*128^ and reference gene embeddings G_e,i_ ∈ ℝ^B*×*128^ was computed as:

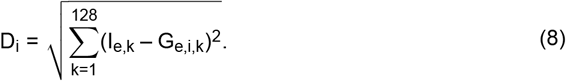

The top 256 closest gene embeddings were selected, and corresponding gene expressions were weighted by a softmax function:

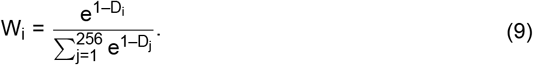

The final retrieved expression profile was the weighted average 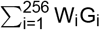.

#### Survival Analysis

Following previous work [54], we employed a deep learning-based approach for survival prediction by discretizing survival time into intervals, with each interval’s survival risk represented by a separate output neuron. This discrete-time approach mitigates the necessity for large batch sizes and facilitates training using individual patient observations. Specifically, we divided the continuous survival time axis into discrete intervals [t_0_, t_1_), [t_1_, t_2_), [t_2_, t_3_), [t_3_, t_4_), with boundaries determined by quartiles of uncensored survival times within each TCGA cohort. Thus, the discrete event time T_j_ for patient j with continuous event time T_j,cont_ is defined as:

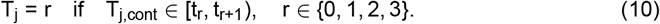

Given discrete-time labels Y_j_ for patient j and their corresponding slide-level feature vector **h**_j,final_, the model’s final layer employs a sigmoid activation function to estimate the hazard function:

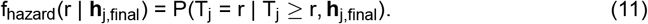

The corresponding survival function, representing the probability that the event time exceeds r, is computed as:

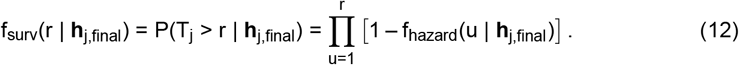

During training, parameters were optimized using the discrete-time log-likelihood loss [**zadeh2020bias**], explicitly accounting for censored observations:

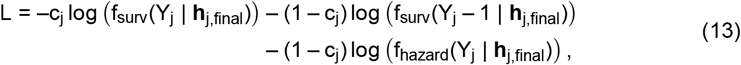

where c_j_ = 1 indicates patient j survives beyond the follow-up period (censored), and c_j_ = 0 indicates an observed event (e.g., death). To further enhance predictive accuracy, we applied a weighted loss emphasizing uncensored cases:

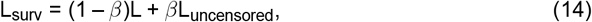

where the uncensored loss term is defined as:

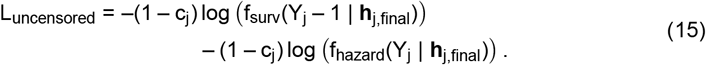

The slide-level feature vector **h**_j,final_ comprises BRIDGE-predicted gene expressions, morphological features extracted via existing pathology foundation models, and patient-wise bulk RNA-seq data. Patch-level morphological features were aggregated into slide-level representations using average and attention-based pooling. For BRIDGE-predicted gene expressions and bulk RNA-seq counts, we selected the top 1,000 highly variable genes (HVGs) as the genomic features.

### Evaluation Metrics

#### Pearson Correlation Coefficient (PCC)

To quantify gene expression prediction accuracy, we computed the Pearson correlation coefficient (PCC) between predicted 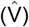 and measured (V) expression levels, defined for a gene across S spatial spots as:

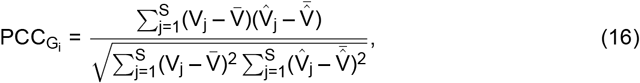

where 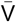 and 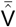 denote means of measured and predicted expression, respectively. Slide-level PCC averages across 80 biomarkers. PCC ranges from –1 (perfect negative correlation) to 1 (perfect positive correlation), with 0 indicating no linear correlation.

#### Concordance Index (C-index)

We evaluated survival analysis using the Concordance index (C-index), quantifying a model’s ability to correctly rank patient survival times, particularly important with censored data:

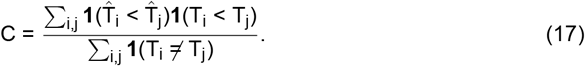

C-index values range from 0.5 (random prediction) to 1.0 (perfect concordance).

## Data Availability

The datasets utilized in this study are broadly categorized into three groups: **(1)** BIG-600K, our self-curated, preprocessed multi-organ spatial transcriptomics dataset; **(2)** External single-cell RNA sequencing (scRNA-seq) datasets, employed specifically for cross-modality retrieval tasks involving single-cell data; **(3)** External cancer cohorts from The Cancer Genome Atlas (TCGA), used for survival analysis evaluations. For certain external single-cell datasets, only cells corresponding explicitly to specific organs were selected to construct organ-specific retrieval pools, in alignment with our experimental design. All datasets used in this study are publicly accessible, with detailed access links and dataset descriptions provided comprehensively in Supplementary Table S4. Additionally, the preprocessed BIG-600K dataset has been made publicly available through Mendeley Data, facilitating straightforward access and reproducibility by the broader scientific community.

## Code Availability

The complete BRIDGE source code developed in this study is publicly available on GitHub (https://github.com/tracy666/BRIDGE). In addition to the source code, we have provided comprehensive documentation, pre-trained model checkpoints, scripts, and detailed instructions necessary to fully reproduce all analyses, experiments, and results presented in this paper. These resources enable researchers and practitioners to readily apply BRIDGE to their own datasets and experimental contexts, facilitating broader adoption and further exploration.

## Acknowledgements

We thank Dr. Lingyu Li (Postdoc of Bioinformatics, LKS Faculty of Medicine, The University of Hong Kong) for insightful discussions and constructive suggestions that improved the manuscript. This work was partially supported by the Research Grants Council of Hong Kong SAR, China (Project No. 27206123 and T45-401/22-N) and the Hong Kong Innovation and Technology Fund (Project No. ITS/274/22).

## Author Contributions

LY and YH conceived, designed, and supervised the study. ZL and WZ developed and implemented the BRIDGE framework, conducted experiments, and performed data analyses, with technical support from FW. GC contributed to data processing and analysis. ZL, WZ, LY, and YH drafted and wrote the manuscript collaboratively, with critical input, discussions, and final approval from all authors.

## Supplemental Figures

**Figure S1.**
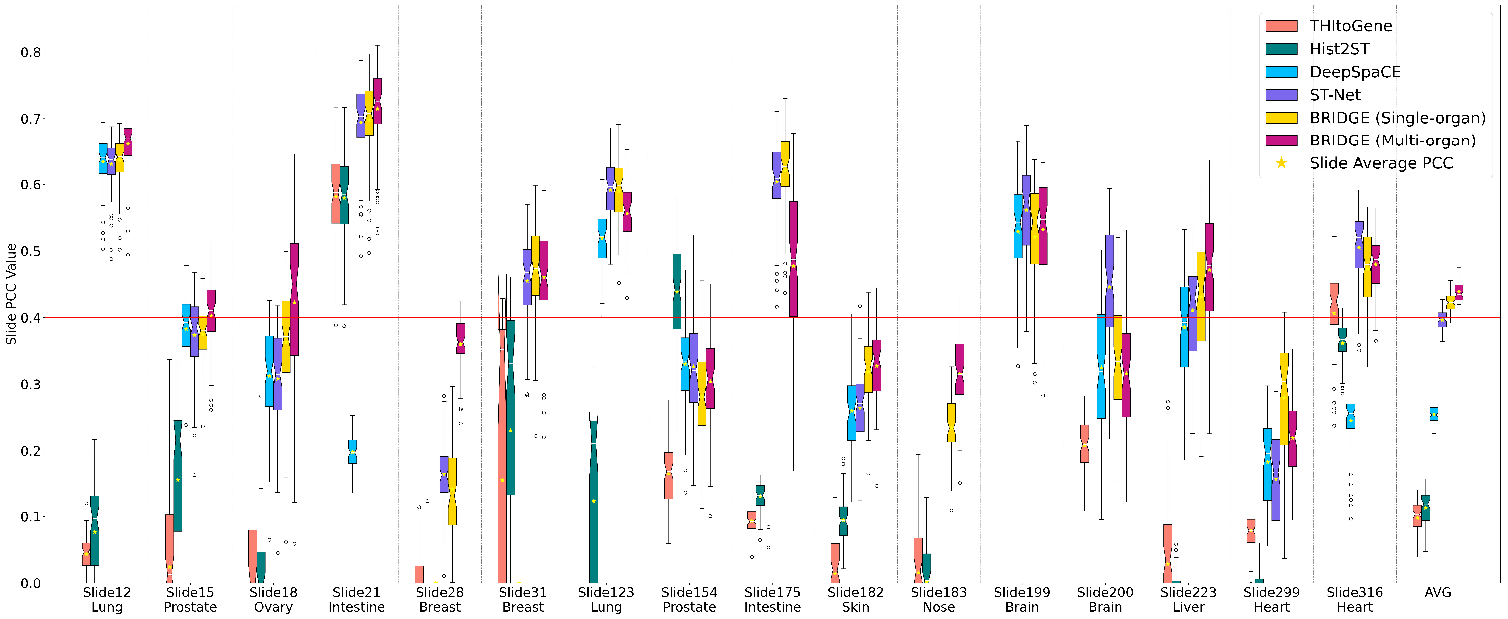
Comprehensive gene expression prediction performance across the multiorgan evaluation set. Detailed histogram illustrating the slide-level Pearson correlation coefficient (PCC) scores for gene expression prediction across all 16 whole-slide images (WSIs) from ten human organs (brain, breast, heart, liver, lung, nasopharynx, ovary, prostate, skin, and intestine) under the multi-organ evaluation setting. The average slide-level PCC across these WSIs is reported as “AVG”. Results demonstrate that the single-organ BRIDGE consistently outperforms the strongest baseline (ST-Net), achieving an average PCC of 0.42 compared to ST-Net’s 0.40. Notably, multi-organ BRIDGE, which leverages heterogeneous pre-training across the entire BIG-600K dataset, exhibits further improved generalization with an average PCC of 0.44, underscoring the benefits of integrative contrastive and generative training across diverse tissues.

**Figure S2.**
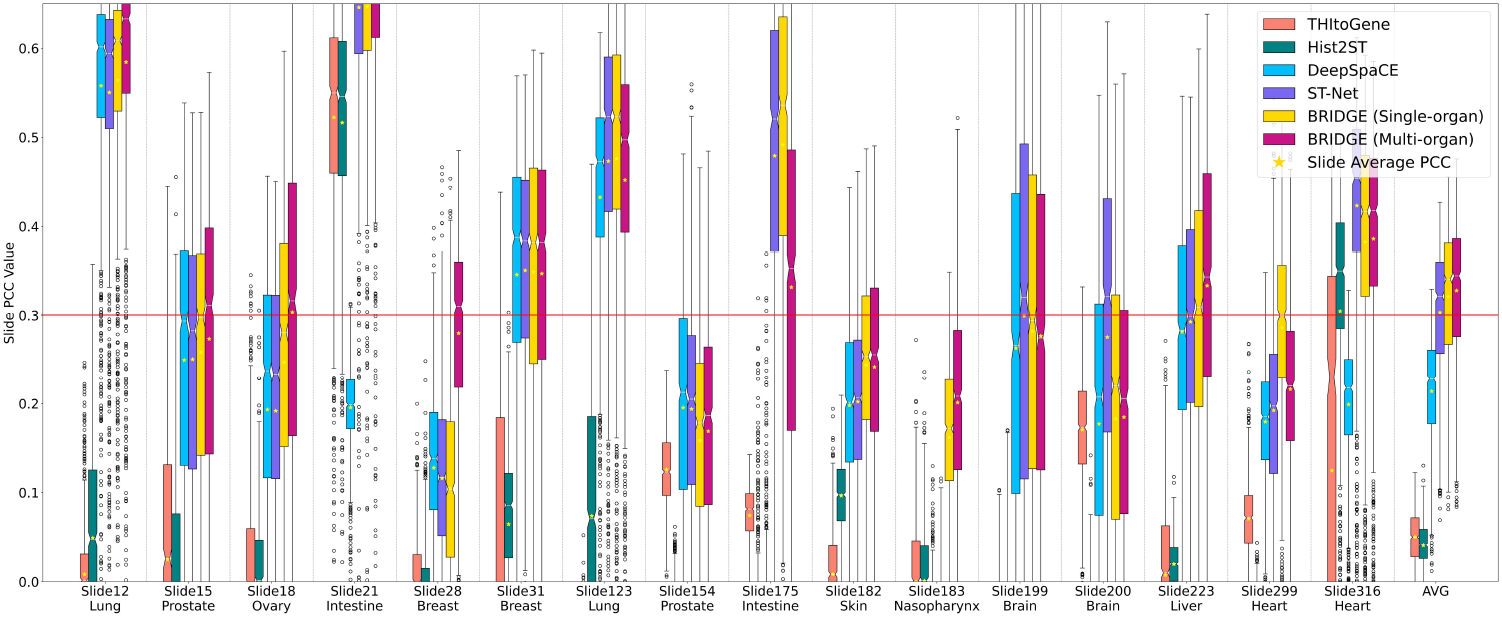
Large-panel gene-expression prediction accuracy across the multi-organ evaluation set. Histograms depict slide-level Pearson correlation coefficients (PCCs) obtained when predicting a 1,056-gene panel—the union of the top-1,000 most highly expressed genes (HEG) and the top-1,000 highly variable genes (HVG)—in 16 whole-slide images (WSIs) from ten human organs. Single-organ BRIDGE outperforms the strongest baseline (ST-Net) with a mean PCC of 0.32 versus 0.30. Multi-organ BRIDGE achieves a further improvement to a mean PCC of 0.33.

**Figure S3.**
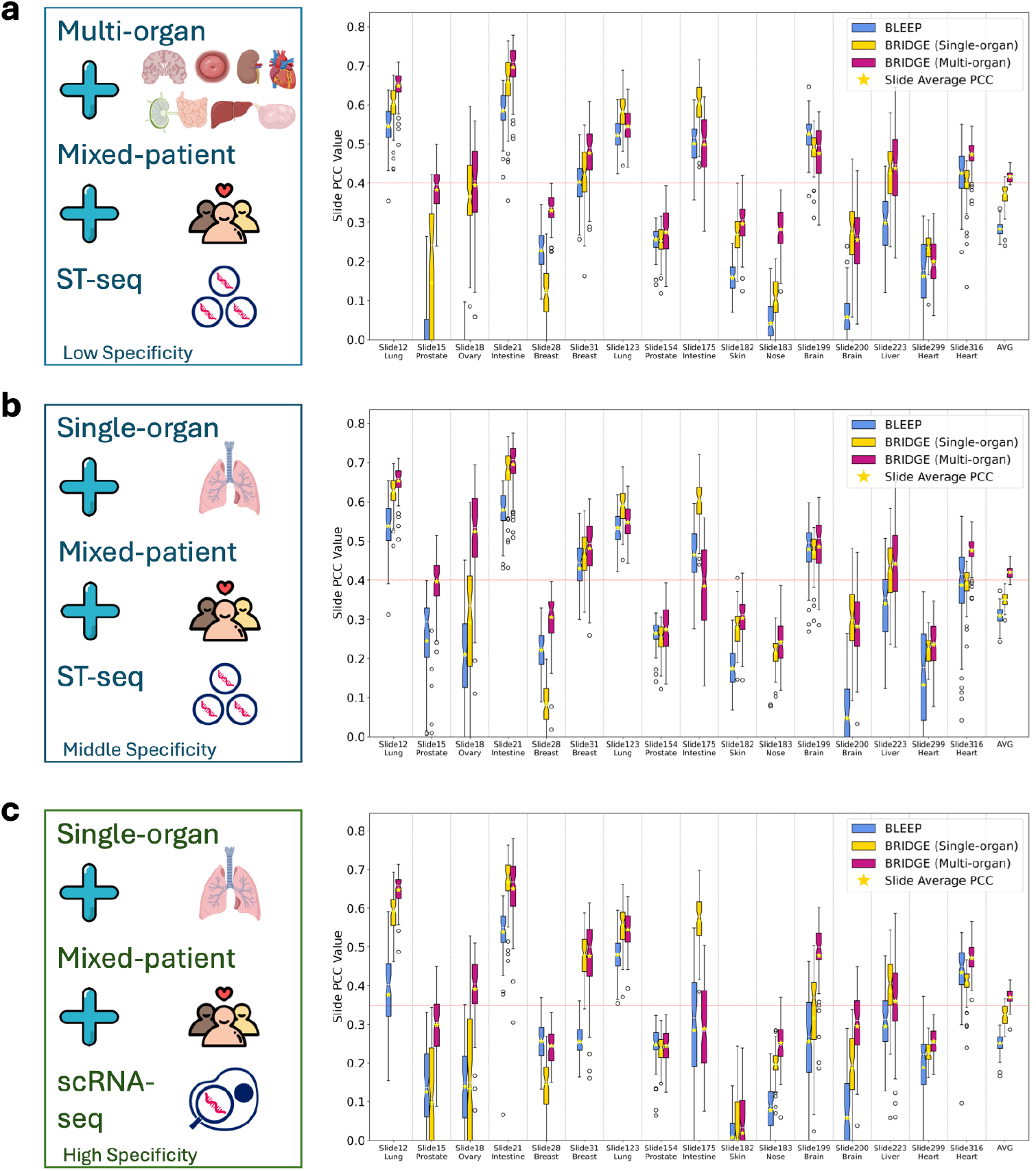
Detailed gene retrieval performance of BRIDGE compared to BLEEP across three gene pool configurations. Histogram showing the slide-level PCC scores for predicting gene expression via retrieval for all 16 WSIs from ten human organs. Performance comparison is conducted across three retrieval pool configurations: (a) Multi-organ ST pool, (b) Single-organ ST pool, and (c) Single-cell RNA-seq (scRNA-seq) pool. BRIDGE consistently achieves higher PCC scores than BLEEP across all retrieval scenarios, highlighting its improved representation learning from multi-organ joint training. Specifically, BRIDGE attains average PCCs of 0.417, 0.420, and 0.370 compared to BLEEP’s 0.283, 0.309, and 0.251 for multi-organ ST, single-organ ST, and scRNA-seq pool configurations, respectively.

**Figure S4.**
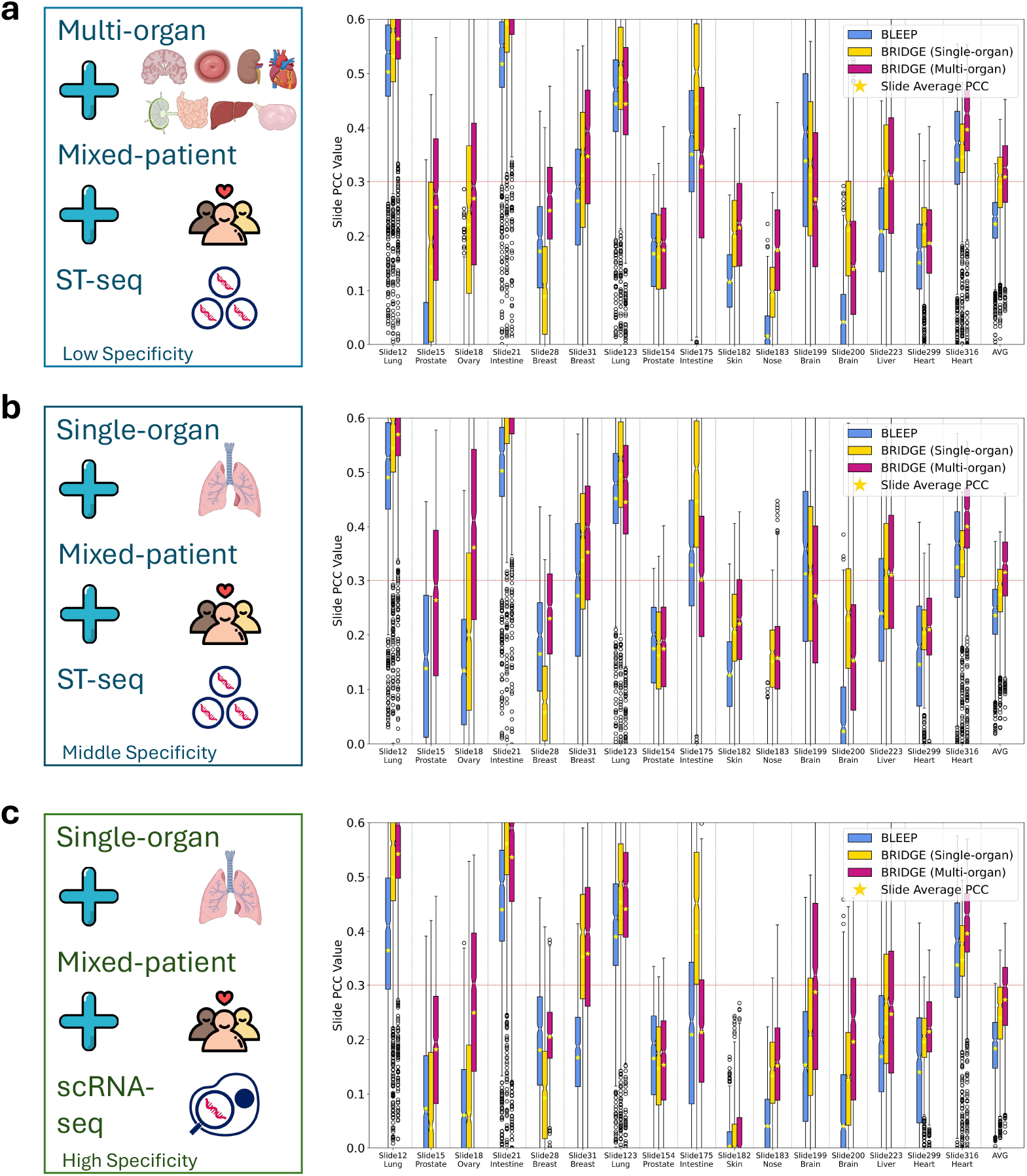
Large-panel gene-expression retrieval performance of BRIDGE versus BLEEP. Histograms display slide-level Pearson correlation coefficients (PCCs) obtained when retrieving a 1,056-gene panel—the union of the 1,000 highly expressed genes (HEG) and the 1,000 highly variable genes (HVG)—from 16 whole-slide images (WSIs) covering ten human organs Across all three settings BRIDGE outperforms the BLEEP baseline, underscoring the benefits of its multi-organ contrastive-and-generative pre-training. Specifically, BRIDGE attains average PCCs of 0.31, 0.32, and 0.27 compared with BLEEP’s 0.22, 0.24, and 0.18 for the multi-organ ST, single-organ ST, and scRNA-seq pools, respectively.

**Figure S5.**
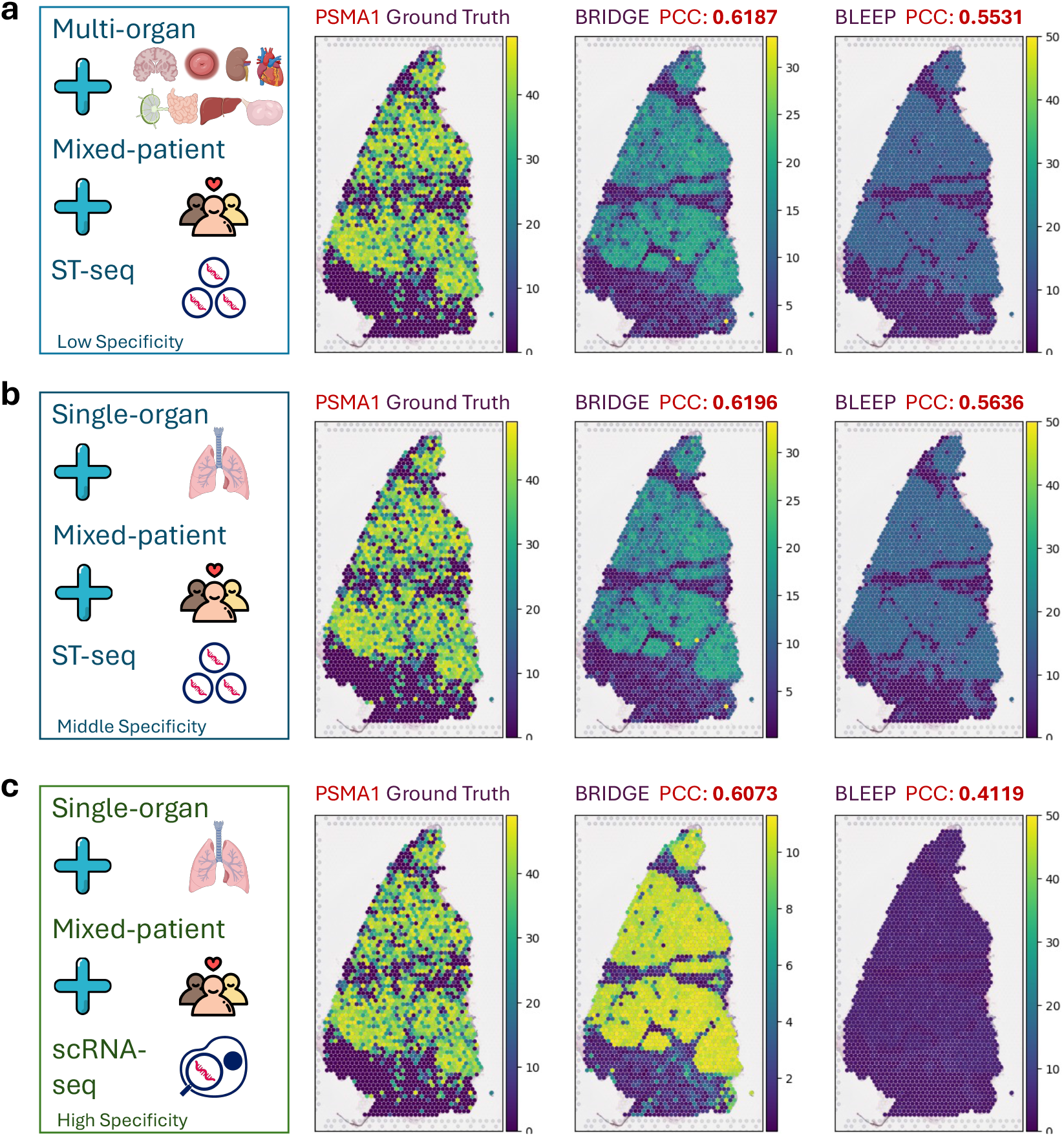
Spatial visualization comparison of retrieved gene expression for lung biomarker gene PSMA1. Spatial expression patterns of lung biomarker gene PSMA1 retrieved by BRIDGE and BLEEP across (a) Multi-organ ST pool, (b) Single-organ ST pool, and (c) Single-cell RNA-seq pool. Visual results illustrate BRIDGE’s superior ability to capture precise and biologically meaningful spatial distributions, consistently surpassing BLEEP in accurately reflecting known spatial gene expression patterns.

**Figure S6.**
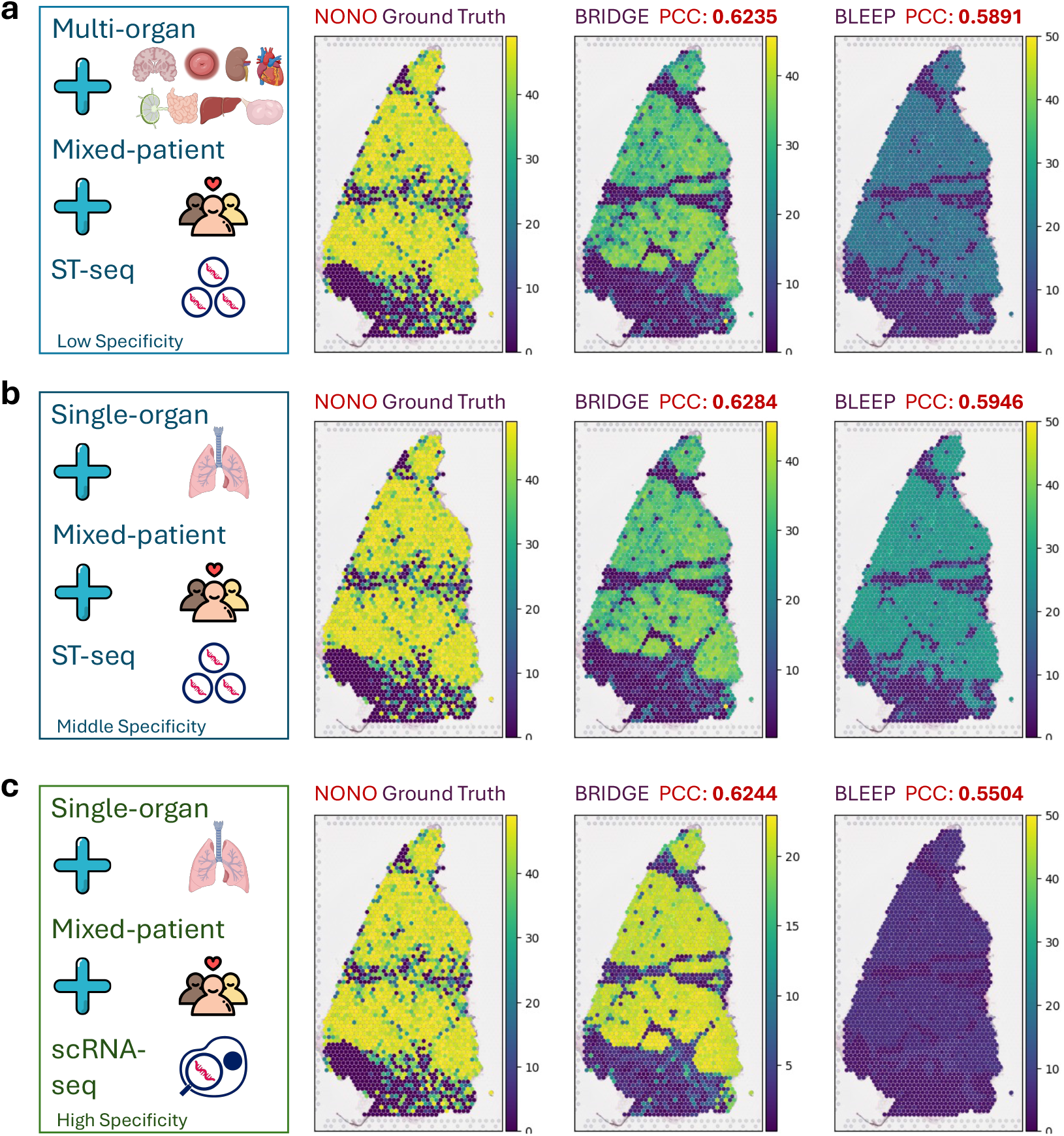
Spatial visualization comparison of retrieved gene expression for lung biomarker gene NONO. Spatially resolved gene expression patterns predicted by BRIDGE and BLEEP for lung biomarker gene NONO, evaluated across three distinct gene retrieval pools: (a) Multi-organ ST pool, (b) Single-organ ST pool, and (c) Single-cell RNA-seq pool. BRIDGE’s retrieval demonstrates more biologically plausible and spatially coherent distribution patterns compared to BLEEP, underscoring BRIDGE’s robust transferability and improved retrieval accuracy across diverse gene pool configurations.

**Figure S7.**
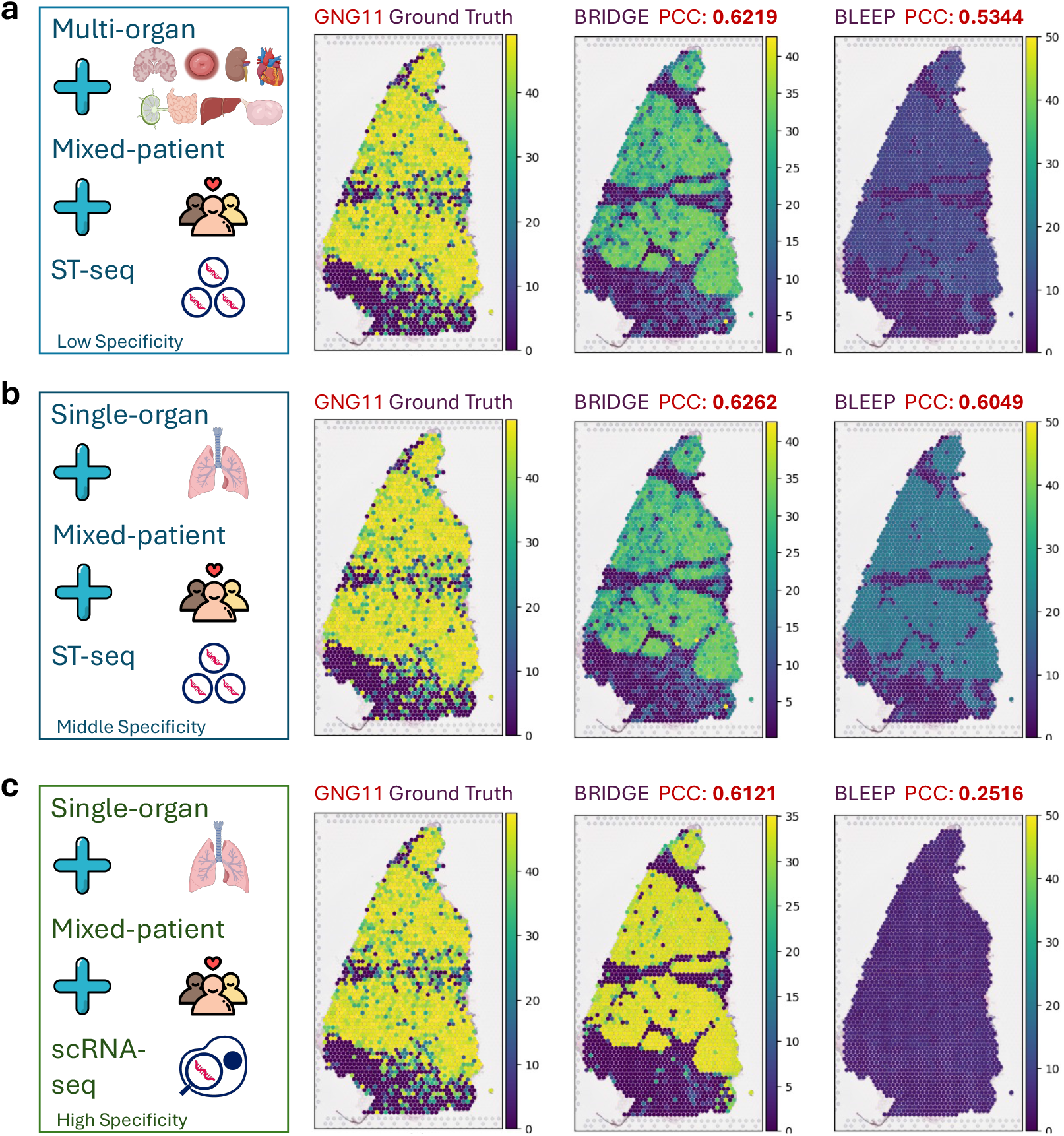
Spatial visualization comparison of retrieved gene expression for lung biomarker gene GNG11. Detailed spatial expression predictions of lung biomarker gene GNG11 using BRIDGE and BLEEP across three retrieval pool settings: (a) Multi-organ ST pool, (b) Single-organ ST pool, and (c) Single-cell RNA-seq pool. BRIDGE consistently provides more accurate and biologically consistent spatial patterns than BLEEP, especially under the modality shift between ST and scRNA-seq, demonstrating the advantage of its integrated contrastive-generative training for effectively transferring knowledge across experimental modalities and conditions.

**Figure S8.**
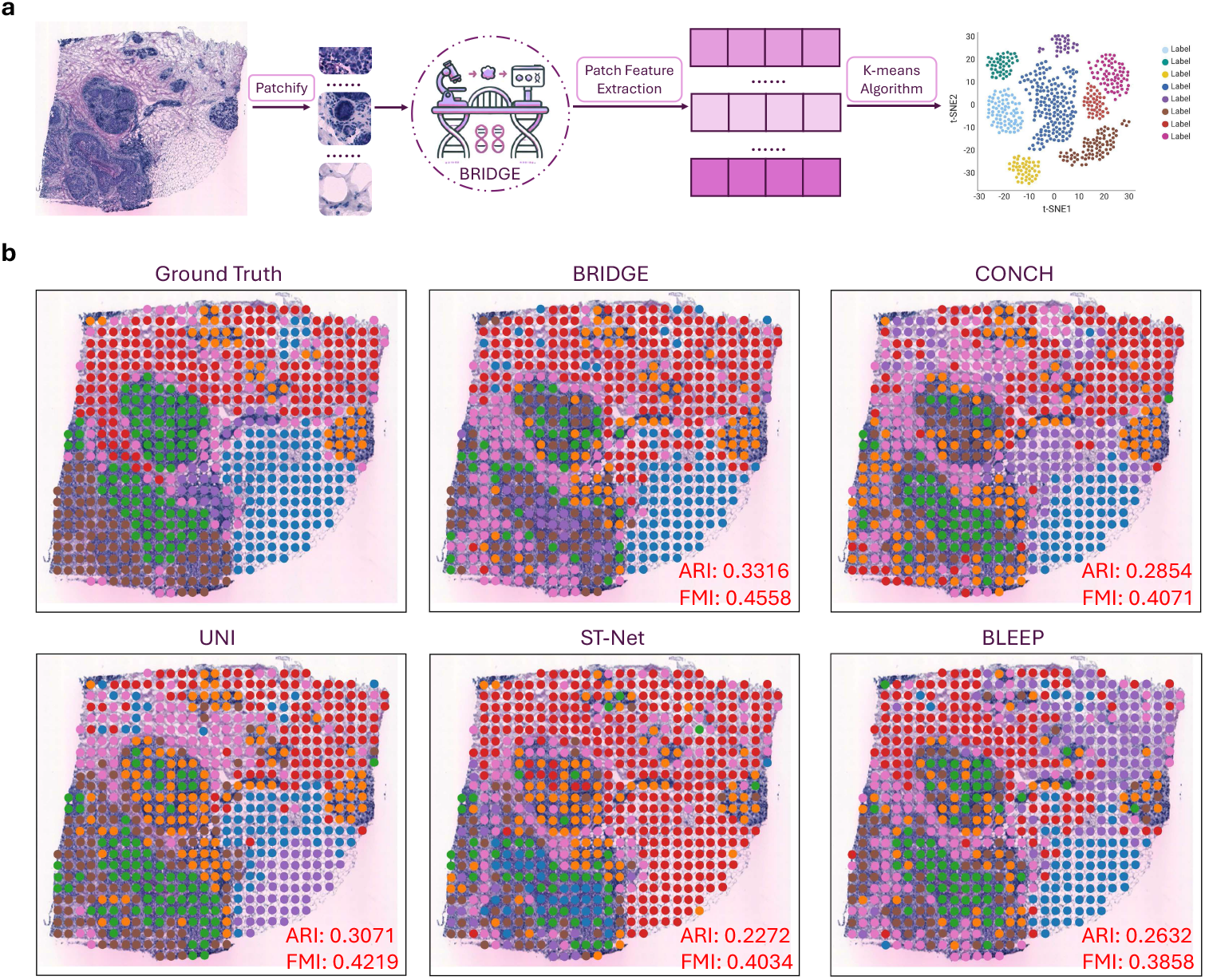
Detailed comparison of cell clustering performance across BRIDGE and alternative models on a pathologist-annotated breast cancer slide. **a**, Overview of the cell clustering pipeline utilized by BRIDGE and comparative models. Initially, the whole-slide image (WSI) is partitioned into smaller image patches. Visual feature representations are subsequently extracted from each patch using the respective image encoders from evaluated models, followed by K-means clustering to assign cell-type clusters to each image patch. **b**, Quantitative and qualitative comparisons of cell clustering results obtained by BRIDGE versus other baseline models, including large-scale pathology foundation models (CONCH [35], UNI [34]) and spatial transcriptomics (ST)-guided models (ST-Net [2], BLEEP [14]). Evaluation is conducted on a human breast cancer WSI expertly annotated by a pathologist into six distinct regions representing diverse tissue types: adipose tissue (blue), breast glands (orange), cancer in situ (green), connective tissue (red), immune infiltrate (light purple), invasive cancer (dark purple), and manually undetermined regions (pink). Patch-level clustering assignments generated by each model are aligned post-hoc to match the expert annotations optimally. Clustering accuracy is quantitatively assessed using Adjusted Rand Index (ARI) and Feature Mutual Information (FMI), two metrics commonly employed to measure clustering consistency and quality. BRIDGE significantly outperforms all baseline models, achieving the highest ARI and FMI scores, indicating its superior capability in accurately delineating distinct cell-type regions. The superior performance of BRIDGE is attributed to its unique integrative framework, which jointly leverages morphological (image-based) and genomic (spatial transcriptomics-based) information through combined contrastive and generative objectives. In contrast, foundation models such as CONCH and UNI, although trained on extensive visual datasets, lack explicit molecular supervision, limiting their ability to resolve subtle molecularly-defined tissue heterogeneity. Similarly, existing ST-guided approaches (ST-Net and BLEEP), despite incorporating genomic data during training, exhibit comparatively lower performance due to less effective integration of multi-modal supervision into their training str_7_ategies.

## Supplemental Tables

**Table S1.**
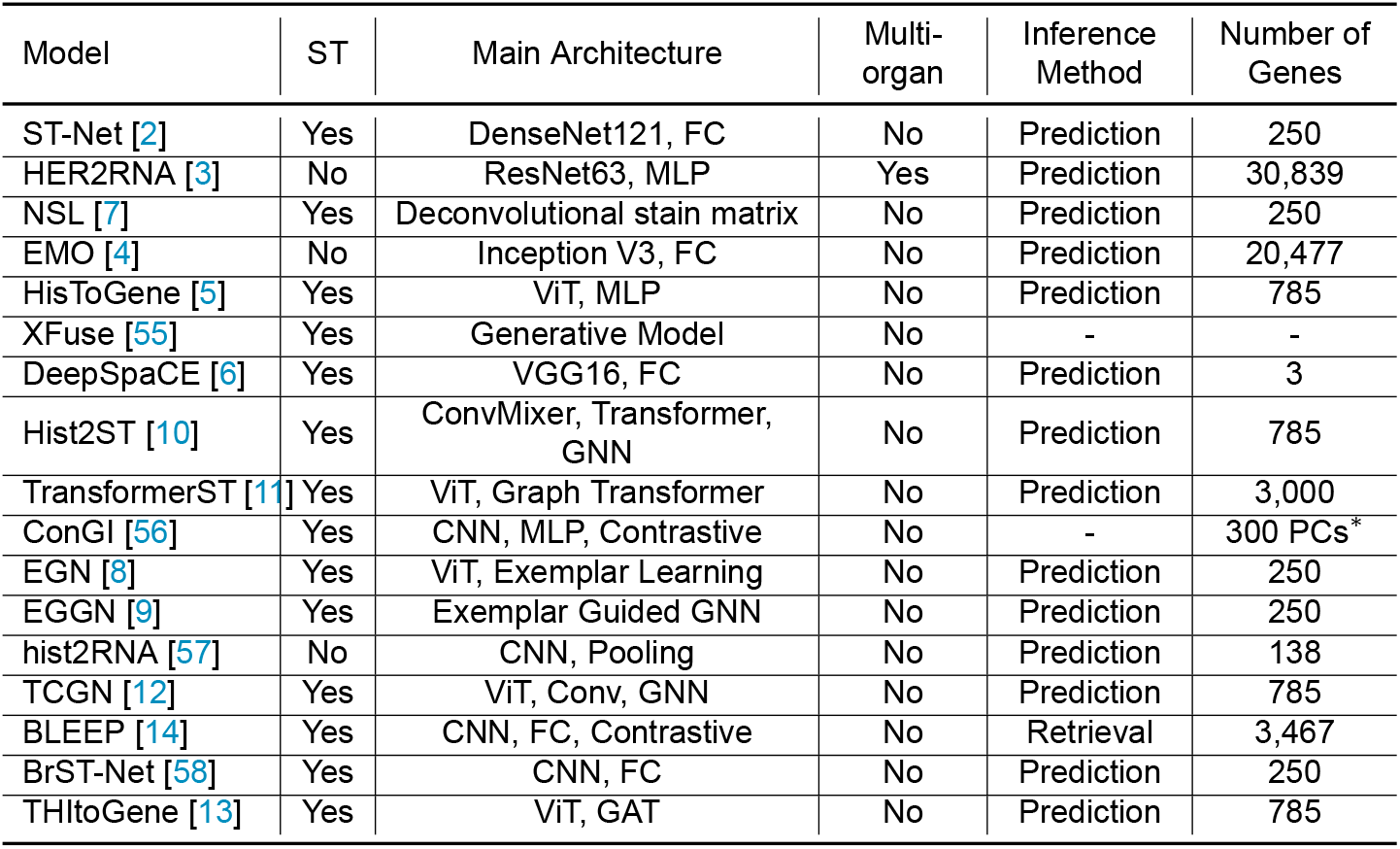
Summary of existing spatial transcriptomics (ST) and bulk RNA-seq prediction models. Comparative summary of prior computational frameworks predicting gene expression or leveraging spatial transcriptomics (ST) and bulk RNA-seq data. The table specifies each model’s architecture, whether it explicitly utilizes ST data, applicability across multiple organs, methods for inferring unobserved gene expression, and the number of genes predicted by each model. Models without explicit use of ST data typically focus on bulk RNA-seq analysis. ^*∗*^PC: Principal Components.

**Table S2.** Numerical composition and summary statistics of the BIG-600K dataset. Detailed numerical summary of the BIG-600K multi-organ spatial transcriptomics dataset, including original dataset sources, sampled organs, number of whole-slide images (WSIs), and total spatial transcriptomic spots. The entries are sorted in descending order by the number of spots.

| Dataset Reference | Organ | Number of Slides | Number of Spots |
| --- | --- | --- | --- |
| 10xGenomics [59] | Breast, Brain, Cervix, Colon, Heart, Intestine, Kidney, Lung, Lymph node, Ovary, Prostate | 28 | 131,263 |
| [60] | Heart | 28 | 97,358 |
| [61] | Brain | 25 | 79,969 |
| [62] | Prostate | 22 | 59,777 |
| [63] | Lung | 15 | 29,863 |
| [64] | Breast | 68 | 30,655 |
| [65] | Lung | 12 | 22,891 |
| [66] | Breast | 36 | 13,620 |
| [67] | Intestine | 4 | 13,119 |
| GSE179572 [68, 69] | Brain | 6 | 12,566 |
| [70] | Skin | 4 | 10,404 |
| GSE240429 [71] | Liver | 4 | 9,296 |
| GSE158328 [72] | Intestine | 8 | 9,330 |
| GSE144239 [73, 74] | Skin | 12 | 8,671 |
| GSE192741 [75, 76] | Liver | 5 | 6,889 |
| [77] | Liver | 4 | 5,993 |
| GSE185715 [78] | Kidney, Lung | 7 | 3,189 |
| GSE171406 [79] | Kidney | 1 | 3,007 |
| GSE200310 [80] | Nasopharynx | 2 | 2,420 |
| GSE178361 [81] | Lung | 2 | 2,220 |

**Table S3.**
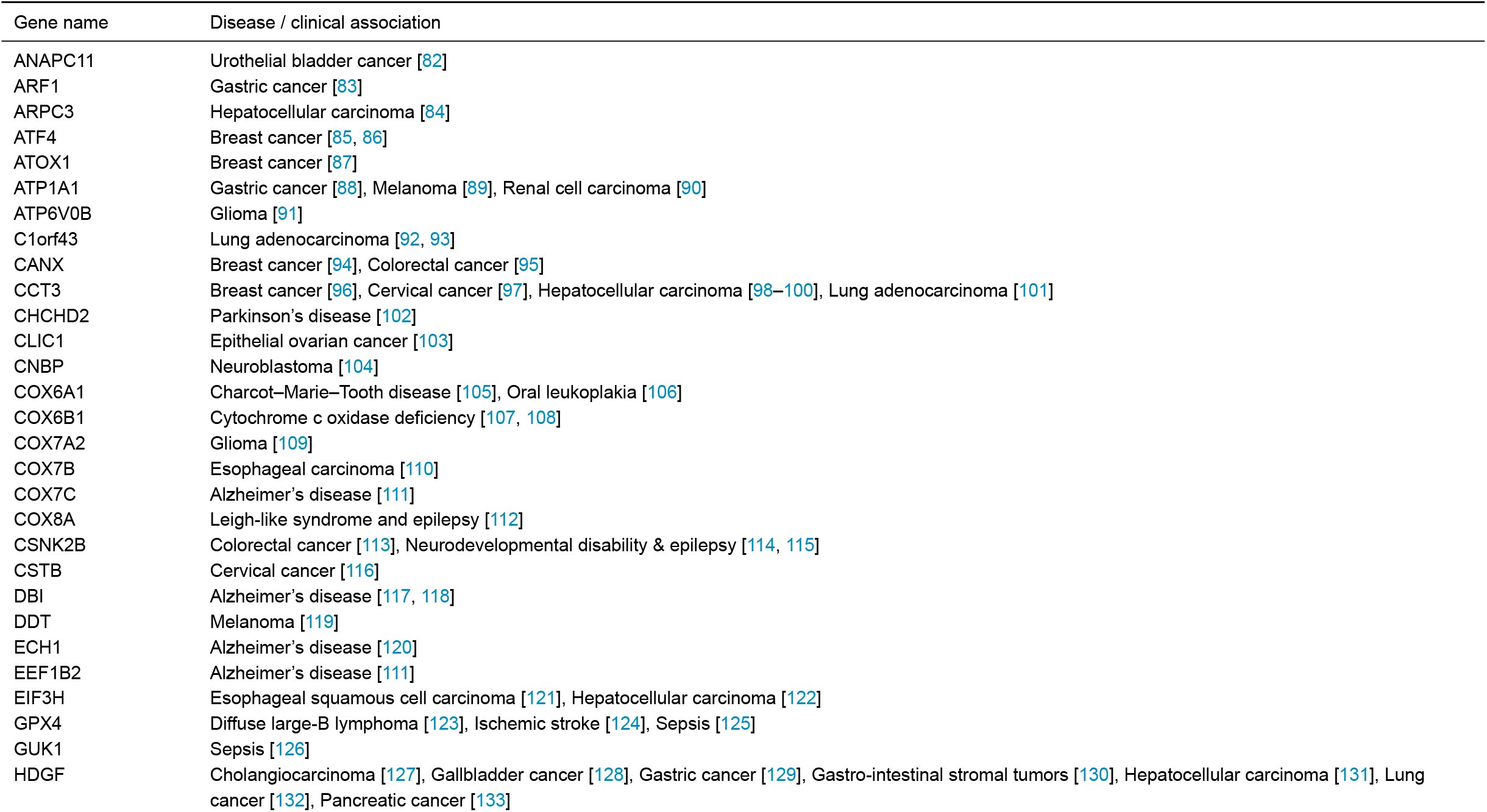
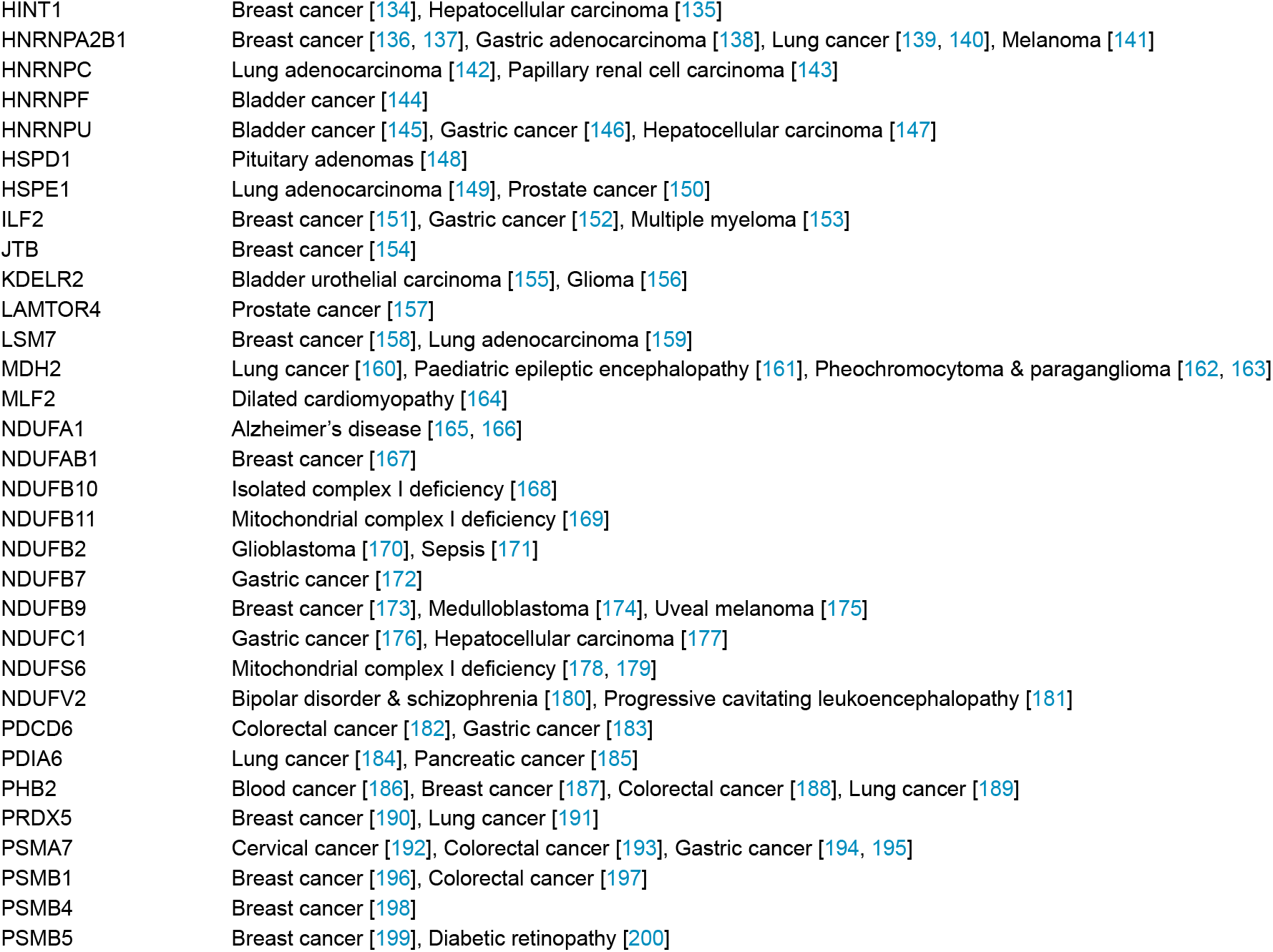
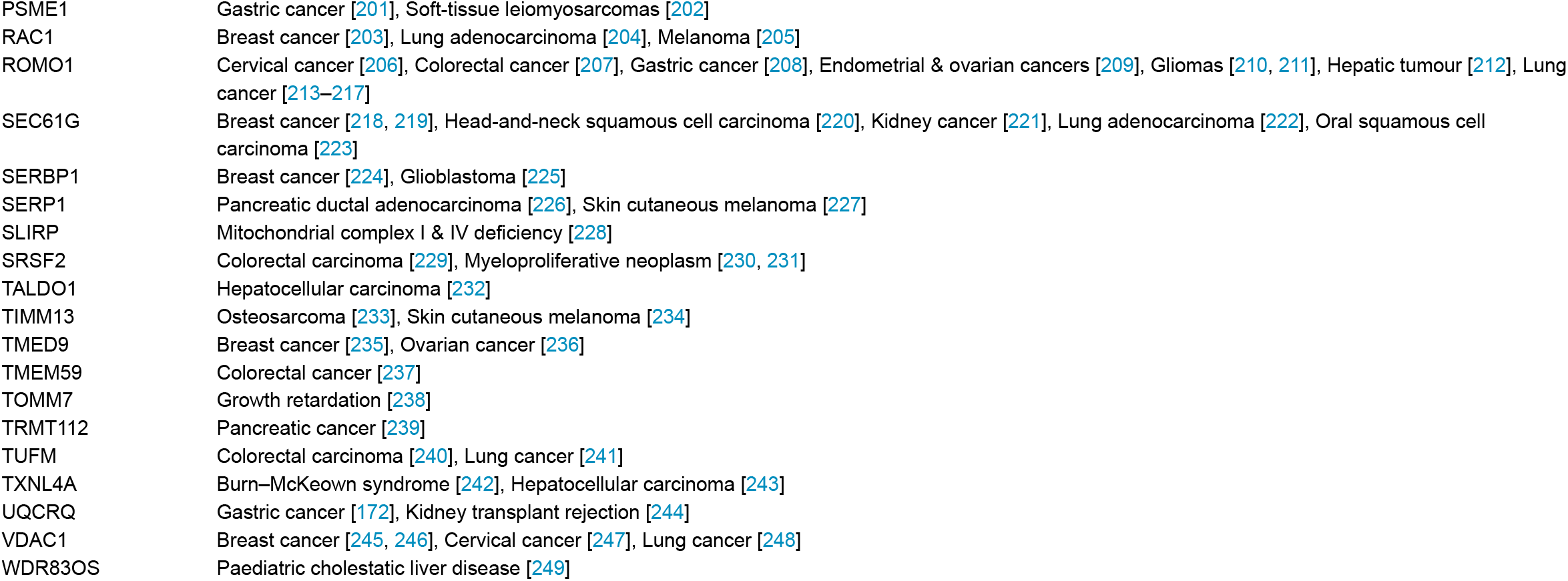
Full list of 80 biomarker genes and their clinical associations. Comprehensive list of the 80 biomarker genes analysed in this study, detailing their known clinical relevance and associations with specific diseases, including various types of cancers and genetic disorders.

| Gene name | Disease / clinical association |
| --- | --- |
| ANAPC11 | Urothelial bladder cancer [82] |
| ARF1 | Gastric cancer [83] |
| ARPC3 | Hepatocellular carcinoma [84] |
| ATF4 | Breast cancer [85, 86] |
| ATOX1 | Breast cancer [87] |
| ATP1A1 | Gastric cancer [88], Melanoma [89], Renal cell carcinoma [90] |
| ATP6V0B | Glioma [91] |
| C1orf43 | Lung adenocarcinoma [92, 93] |
| CANX | Breast cancer [94], Colorectal cancer [95] |
| CCT3 | Breast cancer [96], Cervical cancer [97], Hepatocellular carcinoma [98–100], Lung adenocarcinoma [101] |
| CHCHD2 | Parkinson's disease [102] |
| CLIC1 | Epithelial ovarian cancer [103] |
| CNBP | Neuroblastoma [104] |
| COX6A1 | Charcot–Marie–Tooth disease [105], Oral leukoplakia [106] |
| COX6B1 | Cytochrome c oxidase deficiency [107, 108] |
| COX7A2 | Glioma [109] |
| COX7B | Esophageal carcinoma [110] |
| COX7C | Alzheimer's disease [111] |
| COX8A | Leigh-like syndrome and epilepsy [112] |
| CSNK2B | Colorectal cancer [113], Neurodevelopmental disability & epilepsy [114, 115] |
| CSTB | Cervical cancer [116] |
| DBI | Alzheimer's disease [117, 118] |
| DDT | Melanoma [119] |
| ECH1 | Alzheimer's disease [120] |
| EEF1B2 | Alzheimer's disease [111] |
| EIF3H | Esophageal squamous cell carcinoma [121], Hepatocellular carcinoma [122] |
| GPX4 | Diffuse large-B lymphoma [123], Ischemic stroke [124], Sepsis [125] |
| GUK1 | Sepsis [126] |
| HDGF | Cholangiocarcinoma [127], Gallbladder cancer [128], Gastric cancer [129], Gastro-intestinal stromal tumors [130], Hepatocellular carcinoma [131], Lung cancer [132], Pancreatic cancer [133] |
| HINT1 | Breast cancer [134], Hepatocellular carcinoma [135] |
| HNRNPA2B1 | Breast cancer [136, 137], Gastric adenocarcinoma [138], Lung cancer [139, 140], Melanoma [141] |
| HNRNPC | Lung adenocarcinoma [142], Papillary renal cell carcinoma [143] |
| HNRNPF | Bladder cancer [144] |
| HNRNPU | Bladder cancer [145], Gastric cancer [146], Hepatocellular carcinoma [147] |
| HSPD1 | Pituitary adenomas [148] |
| HSPE1 | Lung adenocarcinoma [149], Prostate cancer [150] |
| ILF2 | Breast cancer [151], Gastric cancer [152], Multiple myeloma [153] |
| JTB | Breast cancer [154] |
| KDELR2 | Bladder urothelial carcinoma [155], Glioma [156] |
| LAMTOR4 | Prostate cancer [157] |
| LSM7 | Breast cancer [158], Lung adenocarcinoma [159] |
| MDH2 | Lung cancer [160], Paediatric epileptic encephalopathy [161], Pheochromocytoma & paraganglioma [162, 163] |
| MLF2 | Dilated cardiomyopathy [164] |
| NDUFA1 | Alzheimer's disease [165, 166] |
| NDUFAB1 | Breast cancer [167] |
| NDUFB10 | Isolated complex I deficiency [168] |
| NDUFB11 | Mitochondrial complex I deficiency [169] |
| NDUFB2 | Glioblastoma [170], Sepsis [171] |
| NDUFB7 | Gastric cancer [172] |
| NDUFB9 | Breast cancer [173], Medulloblastoma [174], Uveal melanoma [175] |
| NDUFC1 | Gastric cancer [176], Hepatocellular carcinoma [177] |
| NDUFS6 | Mitochondrial complex I deficiency [178, 179] |
| NDUFV2 | Bipolar disorder & schizophrenia [180], Progressive cavitating leukoencephalopathy [181] |
| PDCD6 | Colorectal cancer [182], Gastric cancer [183] |
| PDIA6 | Lung cancer [184], Pancreatic cancer [185] |
| PHB2 | Blood cancer [186], Breast cancer [187], Colorectal cancer [188], Lung cancer [189] |
| PRDX5 | Breast cancer [190], Lung cancer [191] |
| PSMA7 | Cervical cancer [192], Colorectal cancer [193], Gastric cancer [194, 195] |
| PSMB1 | Breast cancer [196], Colorectal cancer [197] |
| PSMB4 | Breast cancer [198] |
| PSMB5 | Breast cancer [199], Diabetic retinopathy [200] |
| PSME1 | Gastric cancer [201], Soft-tissue leiomyosarcomas [202] |
| RAC1 | Breast cancer [203], Lung adenocarcinoma [204], Melanoma [205] |
| ROMO1 | Cervical cancer [206], Colorectal cancer [207], Gastric cancer [208], Endometrial & ovarian cancers [209], Gliomas [210, 211], Hepatic tumour [212], Lung cancer [213–217] |
| SEC61G | Breast cancer [218, 219], Head-and-neck squamous cell carcinoma [220], Kidney cancer [221], Lung adenocarcinoma [222], Oral squamous cell carcinoma [223] |
| SERBP1 | Breast cancer [224], Glioblastoma [225] |
| SERP1 | Pancreatic ductal adenocarcinoma [226], Skin cutaneous melanoma [227] |
| SLIRP | Mitochondrial complex I & IV deficiency [228] |
| SRSF2 | Colorectal carcinoma [229], Myeloproliferative neoplasm [230, 231] |
| TALDO1 | Hepatocellular carcinoma [232] |
| TIMM13 | Osteosarcoma [233], Skin cutaneous melanoma [234] |
| TMED9 | Breast cancer [235], Ovarian cancer [236] |
| TMEM59 | Colorectal cancer [237] |
| TOMM7 | Growth retardation [238] |
| TRMT112 | Pancreatic cancer [239] |
| TUFM | Colorectal carcinoma [240], Lung cancer [241] |
| TXNL4A | Burn–McKeown syndrome [242], Hepatocellular carcinoma [243] |
| UQCRQ | Gastric cancer [172], Kidney transplant rejection [244] |
| VDAC1 | Breast cancer [245, 246], Cervical cancer [247], Lung cancer [248] |
| WDR83OS | Paediatric cholestatic liver disease [249] |

**Table S4.**
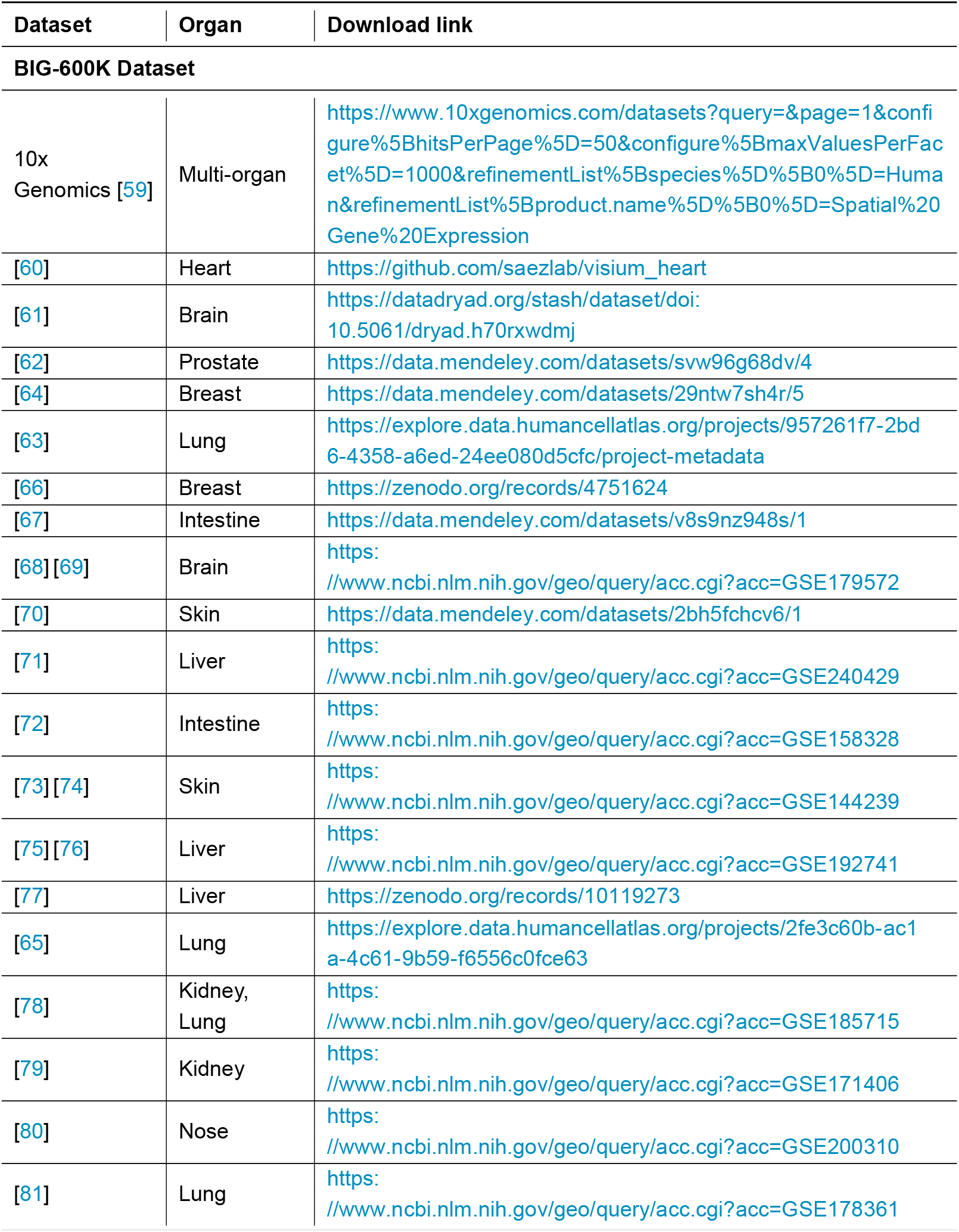
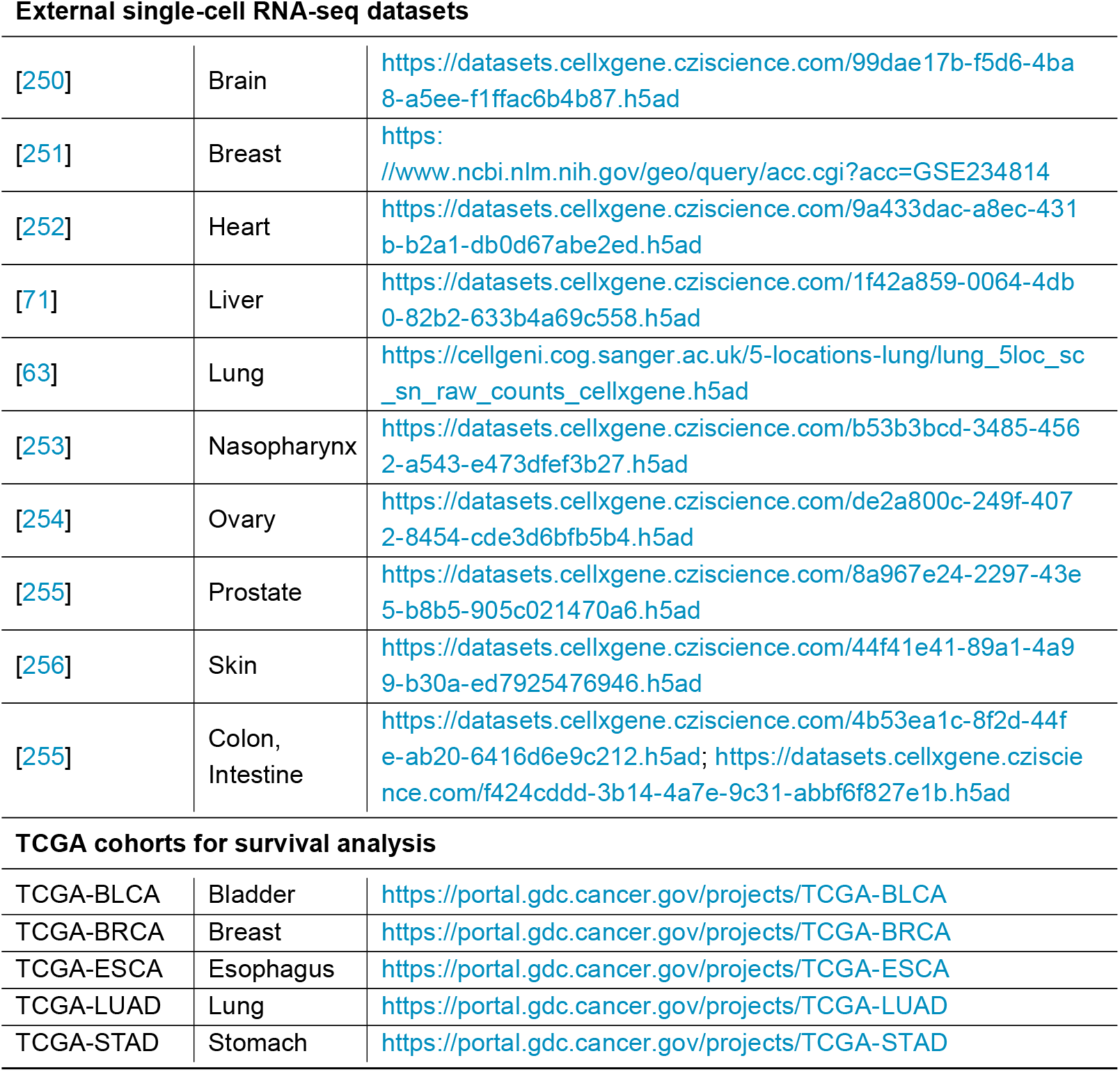
Comprehensive dataset availability and download links for all datasets utilised in this study. This table provides direct access links to (1) the BIG-600K multi-organ spatial-transcriptomics dataset, (2) external single-cell RNA-seq datasets used for retrieval analysis, and (3) TCGA cancer cohorts employed for survival analysis.

| Dataset | Organ | Download link |
| --- | --- | --- |
| **BIG-600K Dataset** |  |  |
| 10x Genomics [59] | Multi-organ | https://www.10xgenomics.com/datasets?query=&page=1&configure%5BhitsPerPage%5D=50&configure%5BmaxValuesPerFacet%5D=1000&refinementList%5Bspecies%5D%5B0%5D=Human&refinementList%5Bproduct.name%5D%5B0%5D=Spatial%20Gene%20Expression |
| [60] | Heart | https://github.com/saezlab/visium_heart |
| [61] | Brain | https://datadryad.org/stash/dataset/doi:10.5061/dryad.h70rxwdmj |
| [62] | Prostate | https://data.mendeley.com/datasets/svw96g68dv/4 |
| [64] | Breast | https://data.mendeley.com/datasets/29ntw7sh4r/5 |
| [63] | Lung | https://explore.data.humancellatlas.org/projects/957261f7-2bd6-4358-a6ed-24ee080d5cfc/project-metadata |
| [66] | Breast | https://zenodo.org/records/4751624 |
| [67] | Intestine | https://data.mendeley.com/datasets/v8s9nz948s/1 |
| [68] [69] | Brain | https://www.ncbi.nlm.nih.gov/geo/query/acc.cgi?acc=GSE179572 |
| [70] | Skin | https://data.mendeley.com/datasets/2bh5fchcv6/1 |
| [71] | Liver | https://www.ncbi.nlm.nih.gov/geo/query/acc.cgi?acc=GSE240429 |
| [72] | Intestine | https://www.ncbi.nlm.nih.gov/geo/query/acc.cgi?acc=GSE158328 |
| [73] [74] | Skin | https://www.ncbi.nlm.nih.gov/geo/query/acc.cgi?acc=GSE144239 |
| [75] [76] | Liver | https://www.ncbi.nlm.nih.gov/geo/query/acc.cgi?acc=GSE192741 |
| [77] | Liver | https://zenodo.org/records/10119273 |
| [65] | Lung | https://explore.data.humancellatlas.org/projects/2fe3c60b-ac1a-4c61-9b59-f6556c0fce63 |
| [78] | Kidney, Lung | https://www.ncbi.nlm.nih.gov/geo/query/acc.cgi?acc=GSE185715 |
| [79] | Kidney | https://www.ncbi.nlm.nih.gov/geo/query/acc.cgi?acc=GSE171406 |
| [80] | Nose | https://www.ncbi.nlm.nih.gov/geo/query/acc.cgi?acc=GSE200310 |
| [81] | Lung | https://www.ncbi.nlm.nih.gov/geo/query/acc.cgi?acc=GSE178361 |
| **External single-cell RNA-seq datasets** |  |  |
| [250] | Brain | https://datasets.cellxgene.cziscience.com/99dae17b-f5d6-4ba8-a5ee-f1ffac6b4b87.h5ad |
| [251] | Breast | https://www.ncbi.nlm.nih.gov/geo/query/acc.cgi?acc=GSE234814 |
| [252] | Heart | https://datasets.cellxgene.cziscience.com/9a433dac-a8ec-431b-b2a1-db0d67abe2ed.h5ad |
| [71] | Liver | https://datasets.cellxgene.cziscience.com/1f42a859-0064-4db0-82b2-633b4a69c558.h5ad |
| [63] | Lung | https://cellgeni.cog.sanger.ac.uk/5-locations-lung/lung_5loc_sc_sn_raw_counts_cellxgene.h5ad |
| [253] | Nasopharynx | https://datasets.cellxgene.cziscience.com/b53b3bcd-3485-4562-a543-e473dfef3b27.h5ad |
| [254] | Ovary | https://datasets.cellxgene.cziscience.com/de2a800c-249f-4072-8454-cde3d6bfb5b4.h5ad |
| [255] | Prostate | https://datasets.cellxgene.cziscience.com/8a967e24-2297-43e5-b8b5-905c021470a6.h5ad |
| [256] | Skin | https://datasets.cellxgene.cziscience.com/44f41e41-89a1-4a99-b30a-ed7925476946.h5ad |
| [255] | Colon, Intestine | https://datasets.cellxgene.cziscience.com/4b53ea1c-8f2d-44fe-ab20-6416d6e9c212.h5ad; https://datasets.cellxgene.cziscience.com/f424cddd-3b14-4a7e-9c31-abbf6f827e1b.h5ad |
| **TCGA cohorts for survival analysis** |  |  |
| TCGA-BLCA | Bladder | https://portal.gdc.cancer.gov/projects/TCGA-BLCA |
| TCGA-BRCA | Breast | https://portal.gdc.cancer.gov/projects/TCGA-BRCA |
| TCGA-ESCA | Esophagus | https://portal.gdc.cancer.gov/projects/TCGA-ESCA |
| TCGA-LUAD | Lung | https://portal.gdc.cancer.gov/projects/TCGA-LUAD |
| TCGA-STAD | Stomach | https://portal.gdc.cancer.gov/projects/TCGA-STAD |

## Footnotes

1 https://www.ensembl.org/biomart/martview/8cc3a4ff5713743f22ba4eaac7846180

